# Predicting Neoantigen Immunogenicity from *In Vivo* Immune Editing

**DOI:** 10.64898/2026.09.17.752425

**Authors:** Timothy J. Sears, Ko-han Lee, María Muñoz Pérez, Robert Rasmussen, Meghana S. Pagadala, Kairi Tanaka, Ajay Subramanian, Everett J. Moding, Maurizio Zanetti, Hannah Carter

**Affiliations:** Bioinformatics and Systems Biology Program, University of California San Diego, La Jolla, CA, USA; Department of Bioengineering, Universidad Carlos III de Madrid, Madrid, Spain; Biomedical Sciences Program, University of California San Diego, La Jolla, CA, USA; School of Biological Sciences, University of California San Diego, La Jolla, CA, USA; Department of Radiation Oncology, Stanford University, Stanford, CA, USA; Stanford Cancer Institute, Stanford University, Stanford, CA, USA; Moores Cancer Center, University of California San Diego, La Jolla, CA, USA; The Laboratory of Immunology, Moores Cancer Center and Department of Medicine, University of California San Diego, La Jolla, CA, USA

**Keywords:** Neoantigens, personalized cancer vaccine, immunotherapy, immunoediting

## Abstract

Neoantigen immunogenicity prediction is fundamental to personalized cancer vaccines, tumor-infiltrating lymphocyte (TIL) therapy, and TCR-T cell engineering. Existing computational predictors rely primarily on in-vitro correlates of peptide presentation or models trained against assay-based reactivity, and they are typically validated within a single therapeutic setting. We reasoned that the most direct evidence of neoantigen immunogenicity is longitudinal in-vivo elimination: under immune checkpoint blockade (ICB), subclones bearing recognized neoantigens are selectively depleted over time. Here, we present the Neoantigen Elimination Model (NEMo), a two-compartment (CD8 and CD4) machine learning classifier trained on the in-vivo editing (IVE) of neoantigens across serially sequenced, ICB-treated tumors. By using mechanistically inspired NeoPrecis features designed to capture determinants of immunogenicity beyond MHC binding affinity, NEMo recovered assay-confirmed immunogenic neoantigens across four independent, unseen clinical settings—pre-existing immunogenicity screening, personalized cancer vaccines, TIL therapy, and a radiotherapy±ICB ctDNA cohort—and stratified progression-free survival more strongly than ELISPOT-confirmed reactivity. The editing signal further revealed an immune-evasion architecture in which oncogenic drivers and neoantigens restricted to lost or silenced HLA alleles are systematically spared from editing.

**Highlights:**

- NEMo learns in-vivo immunoediting to prioritize immunogenic targets
- NEMo outperforms existing tools across four independent clinical settings
- Complete subclonal coverage and presentation integrity predict survival
- Clonal oncogenic driver status and HLA damage spare neoantigens from editing

## Introduction

Cancer treatment is moving decisively toward personalized immunotherapies that mobilize or expand a patient’s own tumor-reactive T cell repertoire, including personalized neoantigen vaccines, adoptive tumor-infiltrating lymphocyte (TIL) therapy, and engineered TCR-T cells[1], [2], [3]. Protein-coding somatic alterations in the tumor genome can give rise to neoantigens—tumor-specific peptides displayed on the cell surface by major histocompatibility complex (MHC) molecules—which serve as targets for each of these immunotherapy strategies. Yet, only a small minority of somatic mutations yield a therapeutically effective target[4]. A viable neoantigen must not only be transcribed, translated, processed and presented by an available MHC at sufficient density, but also recognized by the patient’s T cell repertoire and remain presented despite tumor evolutionary dynamics[5]. A tumor can carry hundreds to thousands of candidate mutations[6], and only a handful meet all these criteria. Computational target prioritization seeks to identify the minority of effective neoantigens, however, false positive rates are high and target selection remains a major bottleneck for personalized immunotherapies[7], [8].

Neoantigen immunogenicity is measured differently across therapeutic settings, and the distinctions matter for how any predictor should be evaluated[9]. Pre-existing (spontaneous) immunity asks whether a patient has already mounted a response to a neoantigen during natural tumor evolution[10], [11]. Vaccine-induced immunity assesses whether exogenous immunization can raise or amplify a response that was previously absent or subclinical[12], [13], [14], [15]. Adoptive-transfer reactivity determines which of a patient’s expanded tumor-reactive T cells actually drive regression; notably, TIL therapy does not select targets by prediction at all, it expands the neoantigen-reactive cells a patient already harbors, and their therapeutic efficacy remains dependent on the antigens they recognize[2], [16]. A model useful across these settings must therefore generalize beyond any single assay definition of immunogenicity.

Computational neoantigen prioritization has advanced substantially. NetMHCpan-family models predict peptide–MHC binding affinity and stability from large-scale eluted ligands, allowing elimination of a majority of peptides that will never be displayed for immune surveillance [17], [18]. As early trials in peptide vaccines have shown, MHC presentation is a necessary prerequisite to immunogenicity, however, it is insufficient to guarantee clinical benefit on its own[19]. Recent methods have made important progress toward predicting immunogenicity by incorporating information beyond antigen presentation: PRIME integrates predicted HLA binding with amino-acid features associated with TCR recognition, using experimentally assessed peptides for training [20]; BigMHC uses deep transfer learning to adapt representations learned from antigen presentation data to immunogenicity prediction [21]; Class-II tools such as TLimmuno2[22] and CD4Episcore[23] extend analogous strategies to MHC-II. However, each o these tools is trained on *in-vitro* or assay-derived reactivity labels — peptide–MHC binding, eluted ligands, or T cell responses measured by functional assays. In a patient, a productive anti-tumor response additionally requires antigen processing and export, dendritic cell uptake and trafficking to draining lymph nodes, priming and clonal expansion, tumor infiltration, and sustained presentation at the tumor site[1], [24]. No existing model is trained on a signal that integrates this entire system in a clinical setting.

In prior work, we quantified the determinants of TCR recognition and assembled a unified feature space spanning presentation binding, foreignness, agretopicity, cross-reactivity, and antigen expression[25]. With those determinants defined, the remaining challenge is identifying a training signal that directly reflects immunogenicity as it occurs in vivo. Immune checkpoint blockade (ICB) exerts T cell-mediated selective pressure on tumors, under which subclones bearing recognized neoantigens are preferentially depleted over time[26]. We reasoned that this longitudinal in-vivo immunoediting signal — which we term in-vivo editing (IVE) — is a direct readout of immunogenicity, and that a model trained on this evolutionary signal would generalize where assay-trained models fail.

We therefore developed the Neoantigen Elimination Model (NEMo), a two-compartment (CD8 and CD4) machine learning model trained on longitudinal in-vivo neoantigen editing across serially sequenced ICB-treated tumors, and asked whether it recovers assay-confirmed immunogenic neoantigens in four independent therapeutic settings in which it was never trained. NEMo identifies immunogenic neoantigens in pre-existing immunogenicity screening, personalized cancer vaccines, TIL therapy, and ctDNA cohorts. It outperforms comparator models in both the pre-existing immunogenicity and cancer-vaccine settings, including methods that explicitly model T-cell recognition. NEMo’s predicted immunogenic neoantigens associate with progression-free survival in cancer vaccine studies and TIL therapy studies where assay-confirmed reactivity does not. Across these analyses, NEMo uncovers an immune-evasion phenomenon in which oncogenic drivers and neoantigens restricted to lost or silenced HLA alleles are spared from editing.

## Results

### An in-vivo neoantigen editing signal in serially sequenced ICB-treated tumors

To capture the selective elimination of tumor subclones under active immunotherapy, we assembled whole-exome sequencing data from 215 patients with paired pre- and post-treatment tumor biopsies, accompanied by tumor RNA sequencing where available (Fig. 1B). The aggregated cohort spanned melanoma (n=98, 46%), a heterogeneous “other” group (n=52, 24%), lung (n=26, 12%), bladder (n=21, 10%), and colorectal (n=18, 8%) cancers.

**Figure 1.**
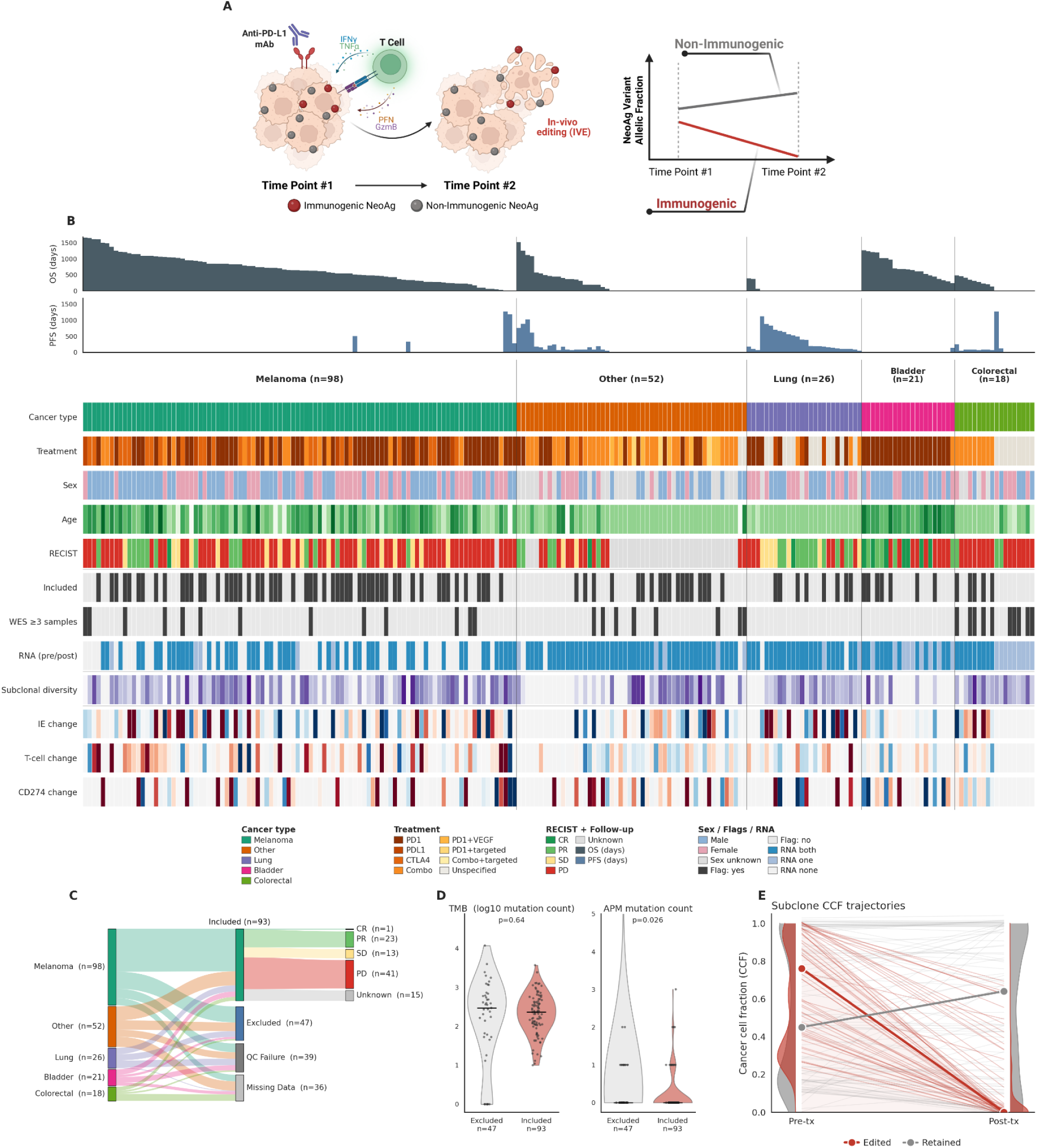
An in-vivo neoantigen editing signal in serially sequenced ICB-treated tumors. **(A)** Study schematic: paired pre- and post-treatment tumor sequencing under ICB is used to track the fate of each neoantigen over therapy, distinguishing eliminated immunogenic mutations (red) from persisting non-immunogenic alterations (grey). **(B)** Oncoprint of 215 paired-timepoint ICB-treated patients, grouped by cancer type. Top bars show overall survival (OS) and progression-free survival (PFS) in days; rows depict per-patient covariates (treatment, RECIST, sex, age, subclonal diversity, immunoediting, T-cell fraction, CD274/PD-L1 expression, and their change between timepoints). **(C)** Flow diagram (Sankey) of cohort selection from initial cancer types (n = 215) through patient inclusion categories (Included, n = 93; Excluded, n = 47; QC Failure, n = 39; and Missing Data, n = 36), alongside RECIST clinical response distribution within the Included subset. **(D)** Violin plots comparing baseline tumor mutational burden (TMB) and antigen presentation machinery (APM) somatic mutation counts between Excluded and Included sets (Mann–Whitney U). **(E)** Per-subclone CCF trajectories from the initial to the extreme post-treatment timepoint, defining IVE-positive (red, collapsing toward zero) versus retained (grey, persisting) mutations; bold lines mark population medians, and marginal curves show CCF density distributions. ICB, immune checkpoint blockade; IVE, in-vivo editing; CCF, cancer cell fraction

Because the in-vivo immunoediting signal we seek to measure relies on the tumor being under active immune pressure, we restricted the analysis to patients with evidence of an ongoing immune response, filtering on immunoediting, T-cell fraction, and CD274 (PD-L1) expression and their change between timepoints. Ninety-three patients met the criteria for active immune engagement (Included), with the remaining tumors being discarded for several reasons (Fig. 1C, Methods). Within the Included set, the full spectrum of clinical outcomes was represented: responders, stable disease, progressive disease and unknowns. Comparing Included and Excluded tumors revealed comparable total mutational burden (TMB P=0.64), but the Excluded group exhibited a significant enrichment in mutations affecting the antigen presentation machinery (APM P=0.026; Fig. 1D), consistent with signs of immune evasion in samples excluded from the analysis. The filter selects on real immune change rather than baseline (Suppl. Fig. 1): overall and progression-free survival did not differ by inclusion status within any cancer type (Suppl. Fig. 2).

To quantify immunoediting, we annotated mutations as to whether they showed evidence of in vivo immune elimination (IVE) across pre- and post-treatment timepoints. Using PhyClone[27], we calculated changes in somatic mutation abundance by measuring the difference in cancer cell fraction (CCF, Methods) from the initial biopsy to the extreme post-treatment timepoint. A mutation was labeled IVE-positive when its CCF declined consistently and substantially across timepoints, defined simultaneously by: an ordinary-least-squares CCF trajectory slope ≤ −0.075, a dynamic range (maximum minus minimum CCF) ≥ 0.25, and an initial CCF ≥ 0.10 (Methods). Throughout, “editing rate” denotes the proportion of mutations in a given stratum that meet this IVE-positive definition. Across the 93 included patients, 22.4% of mutations were IVE-positive (Fig. 1E), with editing rates above 5% observed across all represented tumor types (Suppl. Fig. 3). Mutations were further annotated with features hypothesized to reflect their immunogenic potential, yielding a dataset of mutation-level predictors and IVE labels for model training.

### NEMo learns an editing signal enriched in subclonal, non-driver neoantigens

NEMo comprises two heads, one per T-cell compartment: a CD8 head operating over MHC class-I peptides and a CD4 head over class-II peptides, each returning a per-mutation probability of in-vivo editing (Methods). Under 10-fold cross-validation, both heads predicted editing above chance expectation, although discrimination was modest (CD8 median AUC 0.537, P = 1.2 × 10⁻⁶; CD4 median AUC 0.531, P = 6.7 × 10⁻⁶; Fig. 2A; Suppl. Fig. 4). Interpretation of this performance is complicated by the structure of the training labels. In-vivo editing labels are defined at the subclonal level, meaning mutations co-occurring within the same subclone track synchronously and are not statistically independent. Consequently, a single immunogenic neoantigen driving elimination of its subclone confers an IVE-positive label on every neutral passenger mutation the subclone carries, whereas an immunogenic neoantigen in a subclone that persists for other reasons—including insufficient T-cell infiltration, HLA damage, or stochastic outgrowth—will be labeled negative. The resulting label set is therefore noisy and influenced by subclonal structure, limiting how directly cross-validated AUC reflects the model’s ability to identify immunogenic neoantigens. Future development of immunoediting trained models will need to overcome these substantial limitations.

**Figure 2.**
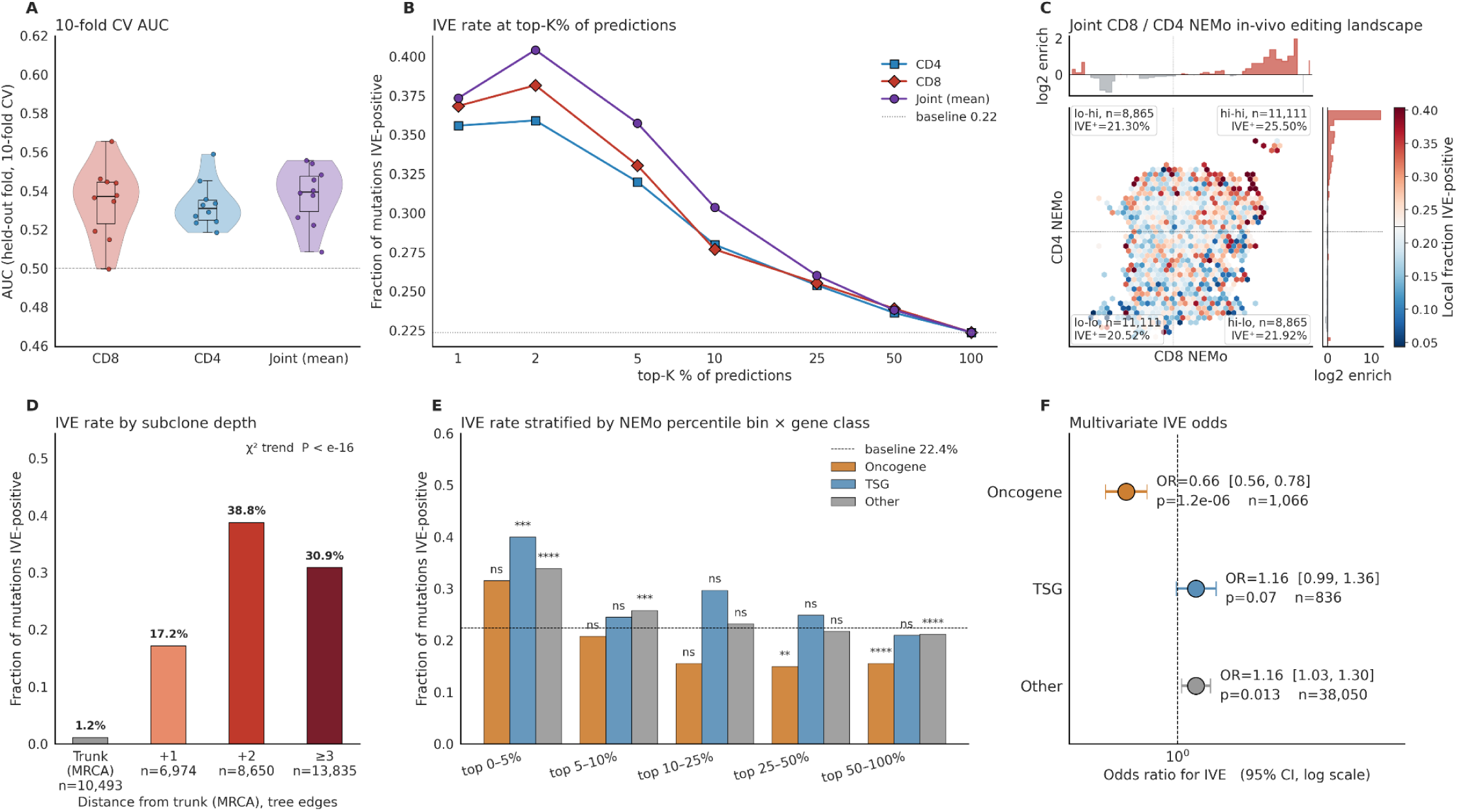
NEMo learns an in-vivo editing signal enriched in subclonal, non-driver neoantigens. **(A)** Held-out AUC across 10 cross-validation folds for the CD8, CD4, and joint heads. **(B)** In-vivo editing (IVE) rate among the top-K% of NEMo predictions relative to the cohort baseline (0.22). **(C)** Joint CD8×CD4 editing landscape (out-of-fold scores) with per-quadrant editing rates. **(D)** IVE rate stratified by subclone tree depth, defined as phylogenetic distance from the trunk/MRCA in the PhyClone tree. **(E)** IVE rate across NEMo prediction percentile bins stratified by canonical gene class (oncogene, TSG, and other) relative to the cohort baseline (22.4%). **(F)** Multivariate forest plot showing adjusted odds ratios (OR, 95% CI) for IVE by gene class from a logistic regression model controlling for NEMo score and cancer cell fraction (CCF). MRCA, most recent common ancestor; TSG, tumor suppressor gene.

The predictive utility of NEMo concentrates on the highest-ranked candidates. IVE rate rose sharply among top-ranked predictions, reaching 0.37–0.40 in the top 1–2% of candidates against a cohort baseline of 0.22 (Fig. 2B). A stepwise comparison on a representative training fold confirmed that incorporating recognition features provides substantial gain over presentation features alone in both the CD8 head (AUC 0.531 vs. 0.554, ΔP = 9.8×10^-8^) and the CD4 head (AUC 0.529 vs. 0.550, ΔP=2.1×10^−10^; Suppl. Fig. 4). Scores from the CD8 and CD4 compartments were only weakly correlated, with the joint model outperforming either alone; neoantigens scored above the median by both heads exhibited an editing rate of 25.5%, compared to 20.5% for those falling below the median in both (Fig. 2C).

We next evaluated how in-vivo editing patterns distribute across tumor evolutionary features. Editing scaled sharply with subclone depth—the phylogenetic distance from the most recent common ancestor (MRCA) in the subclonal tree. Truncal mutations at the MRCA had an editing rate of 1.1%, rising to 17.2% at one node from the trunk, 38.8% at two nodes, and 30.9% at three or more nodes (χ²-trend P << 0.0001; Fig. 2D). Somatic alterations acquired later in tumor evolution were thus substantially more vulnerable to IVE than truncal mutations, consistent with immunogenic potential accumulating alongside additional mutations [28].

We next evaluated IVE by gene category, distinguishing oncogenes, tumor suppressor genes, and genes not assigned to either category (Methods). Across all score bins, mutations in oncogenes exhibited consistently lower editing rates than mutations in either of the other categories (Fig. 2E). In unadjusted univariate analyses, oncogenes showed significantly lower editing rates (16.2% vs. 22.4% baseline; OR=0.67, P=4.7×10^−7^), whereas tumor suppressors (25.1%; OR=1.17, P=0.059) and other genes (22.5%; OR=1.15, P=0.015) tracked above baseline (Suppl. Fig. 5). This depletion held after adjustment for NEMo score and CCF in a multivariate logistic regressor (oncogene OR 0.66, 95% CI 0.56–0.78, P=1.2×10⁻⁶; Fig. 2F). Both tumor suppressors and unclassified categories trended toward higher editing, although tumor suppressors did not reach statistical significance individually (TSG OR 1.16, P=0.07; other OR 1.16, P=0.013; Fig. 2F), and we therefore emphasize the specific depletion of IVE for oncogenic drivers rather than a graded hierarchy across gene classes. Truncal oncogenes represented the least frequently edited class of somatic alterations across the cohort.

This evolutionary distribution could reflect an immune survivorship dynamic wherein tumors become clinically established only if their founding clonal drivers were non-immunogenic or achieved immune escape. In agreement with this model, the editing observed under ICB acts on the subclonal, non-driver periphery of an already established tumor.

### NEMo recovers pre-existing immunogenic neoantigens in independent screening cohorts

We first evaluated whether a model trained on ICB-driven editing could recover neoantigens independently confirmed as immunogenic by functional assay. We repeated this analysis in two validation settings. First, in the NCI Parkhurst cohort, candidate neoantigens from patient tumors were synthesized and screened against autologous T cells for pre-existing reactivity, yielding confirmed immunogenic peptides alongside a well-controlled set of screened-negative peptides within the same patients[10]. In the Wells cohort (the TESLA consortium challenge), participating teams submitted ranked neoantigen predictions for shared patients, and a common set of candidates was assessed for T cell recognition using barcoded peptide–MHC multimers on patient-derived samples[11]. Both cohorts therefore measure pre-existing immunity: whether T cells specific to a given neoantigen are already present in the patient repertoire. Because both platforms supply peptides exogenously—loaded onto recombinant multimers or synthetic reagents—the tumor’s own antigen-presentation status does not influence the readout. Together the two cohorts comprise 83 patients, over 5,000 neoantigens, and 147 confirmed immunogenic peptides (Fig. 3A).

**Figure 3.**
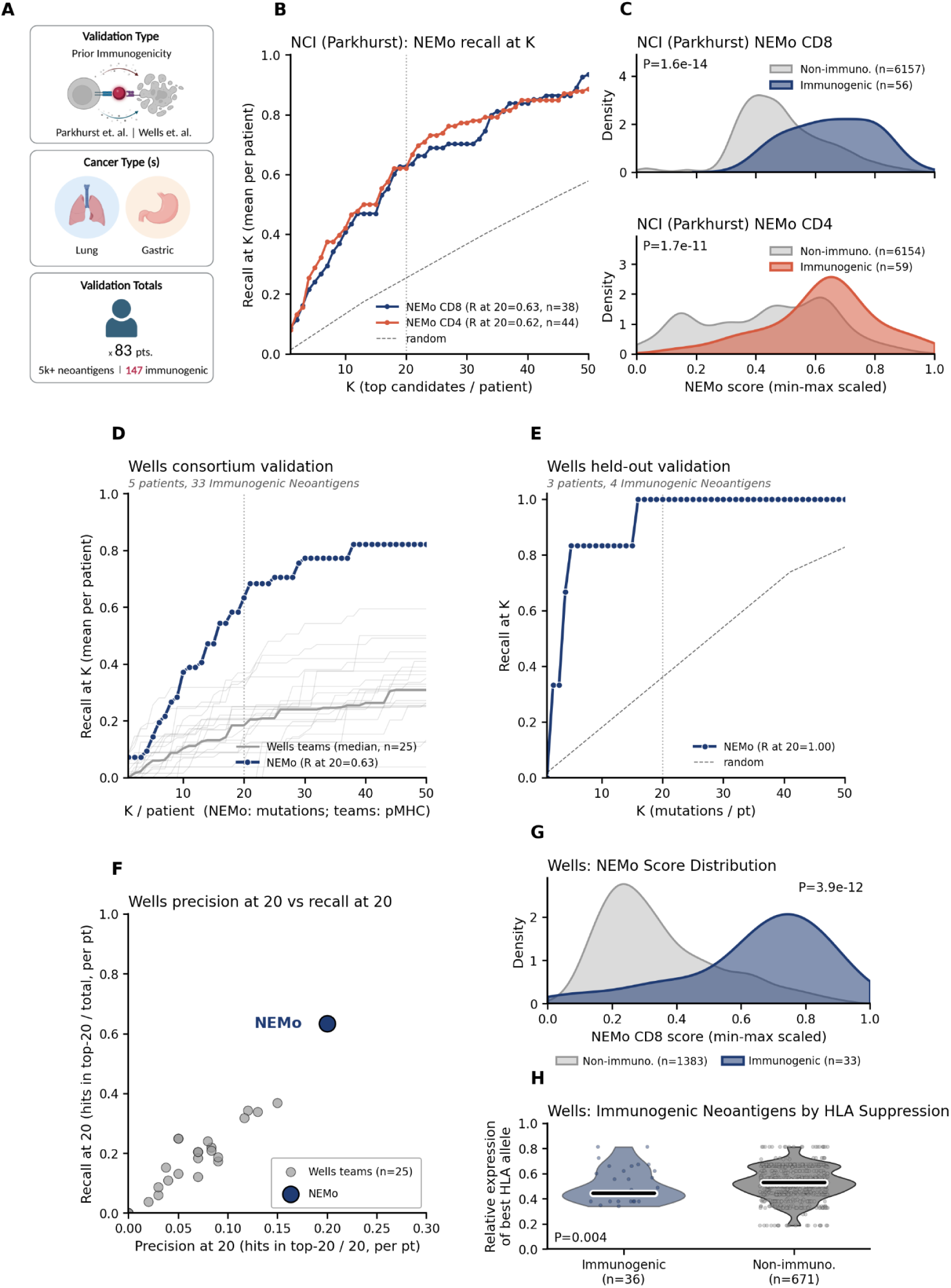
NEMo recovers pre-existing immunogenic neoantigens in independent screening cohorts. Validation in the NCI Parkhurst screening cohort and the Wells cohort (TESLA consortium). **(A)** Schematic of the two cohorts and their cancer types. **(B)** Per-patient recall-at-K of confirmed immunogenic neoantigens in the Parkhurst cohort for the NEMo CD8 and CD4 heads compared to a random baseline. **(C)** Min–max scaled NEMo score distributions for confirmed immunogenic hits versus screened-negative peptides in Parkhurst for CD8 and CD4 (Mann–Whitney U). **(D)** Recall-at-K in the five-patient Wells main challenge analysis, comparing NEMo versus competing consortium teams. **(E)** Recall-at-K in the three held-out Wells patients, evaluated against author-scored ground truth. **(F)** Precision-at-20 versus recall-at-20 for NEMo relative to the competing-team cloud. **(G)** NEMo score separation between confirmed immunogenic and non-immunogenic peptides in the Wells cohort. **(H)** Relative expression of the presenting HLA allele in patient tumors for confirmed immunogenic vs non-immunogenic neoantigens in the Wells cohort (Mann-Whitney U).

In the Parkhurst cohort, NEMo recovered around 63% of a patient’s immunogenic neoantigens within their top 20 ranked candidates in both compartments (CD8 recall@20 = 0.63, n = 38; CD4 recall@20 = 0.62, n = 44; Fig. 3B), and confirmed hits were significantly separated from screened negatives (CD8 P = 1.6×10^-14^; CD4 P = 1.7×10^-11^; Fig. 3C). We selected a cutoff of K=20 for comparisons across analyses because it corresponds to the typical peptide capacity of personalized vaccine formulations[29] (Methods). Benchmarked against established models across Parkhurst patients, NEMo outperformed standard MHC-binding predictors and published immunogenicity algorithms, achieving a relative 21.2% recall advantage in CD8 and a 12.3% advantage in CD4 at K=20 over the next-best comparator (Suppl. Fig. 6A–D). Pairwise score correlations confirmed that NEMo captures distinct predictive features, exhibiting only moderate correlation with presentation-only tools (Suppl. Fig. 6E,F).

The Wells cohort provides a head-to-head comparison, with 25 independent teams predicting on shared patients. In the five-patient main analysis (33 immunogenic mutations), NEMo’s recall-at-K curve exceeded every competing team across the full range of K (NEMo recall@20 = 0.63 versus a team median of ∼0.2; Fig. 3D). In the three held-out validation patients (4 immunogenic mutations), scored exclusively by the study authors, NEMo captured every confirmed immunogenic mutation within the top 15 candidates (recall@20 = 1.00; Fig. 3E). Jointly on precision and recall at 20, NEMo sat outside the competing-team cloud (precision 0.20 vs. team median 0.075; recall 0.63 vs. 0.2; Fig. 3F), scoring immunogenic peptides significantly higher than non-immunogenic ones (P = 3.9×10⁻^12^; Fig. 3G).

When evaluated against standalone algorithms across all Wells patients (n=8 patients, 5 main consortium pts., 3 validation, 37 immunogenic targets), NEMo achieved a recall@20 of 0.77, substantially outperforming competing methods, ranging from recall@20 of 0.47-0.63 (Suppl. Fig. 7A). This represented a 21.98% relative recall advantage over the next-best tool at the clinically relevant K=20 threshold (Suppl. Fig. 7B), while correlating only moderately with conventional affinity predictors (Suppl. Fig. 7C).

Finally, we examined the antigen presentation context in the Wells cohort, which contained no confirmed HLA loss-of-heterozygosity or loss-of-function events, restricting the analysis to allele-specific expression. Immunogenic Wells peptides were preferentially presented by HLA-I alleles that were allele-specifically down-regulated in the tumor (Mann-Whitney U P = 0.004; Fig. 3H). Because multimer assays supply MHC exogenously, they cannot reflect an effect of tumor-specific HLA expression on detection. Instead, this observation suggests that alleles responsible for presenting immunogenic neoantigens could be selectively repressed during tumor evolution in an immunocompetent microenvironment.

### NEMo-predicted neoantigens track outcome in personalized cancer vaccine cohorts

We next applied NEMo to personalized neoantigen cancer vaccine cohorts, wherein identifying immunogenic neoantigens is the explicit intent of target selection and targets can be linked to clinical outcome. In these trials (Cafri, Ott, Blass, and Braun[12], [13], [14], [15]) patients received synthetic long peptides with adjuvant, and immunogenicity was determined after vaccination by IFN-γ ELISPOT on peripheral blood mononuclear cells, with intracellular cytokine staining used in some studies to attribute responses to CD4 or CD8 compartments[12], [13], [14], [15]. Truth labels here therefore denote vaccine-induced reactivity — whether immunization successfully raised or amplified a detectable T-cell response. Together the four cohorts comprise 30 patients, 358 screened neoantigens, and 149 assay-confirmed immunogenic peptides (Fig. 4A).

**Figure 4.**
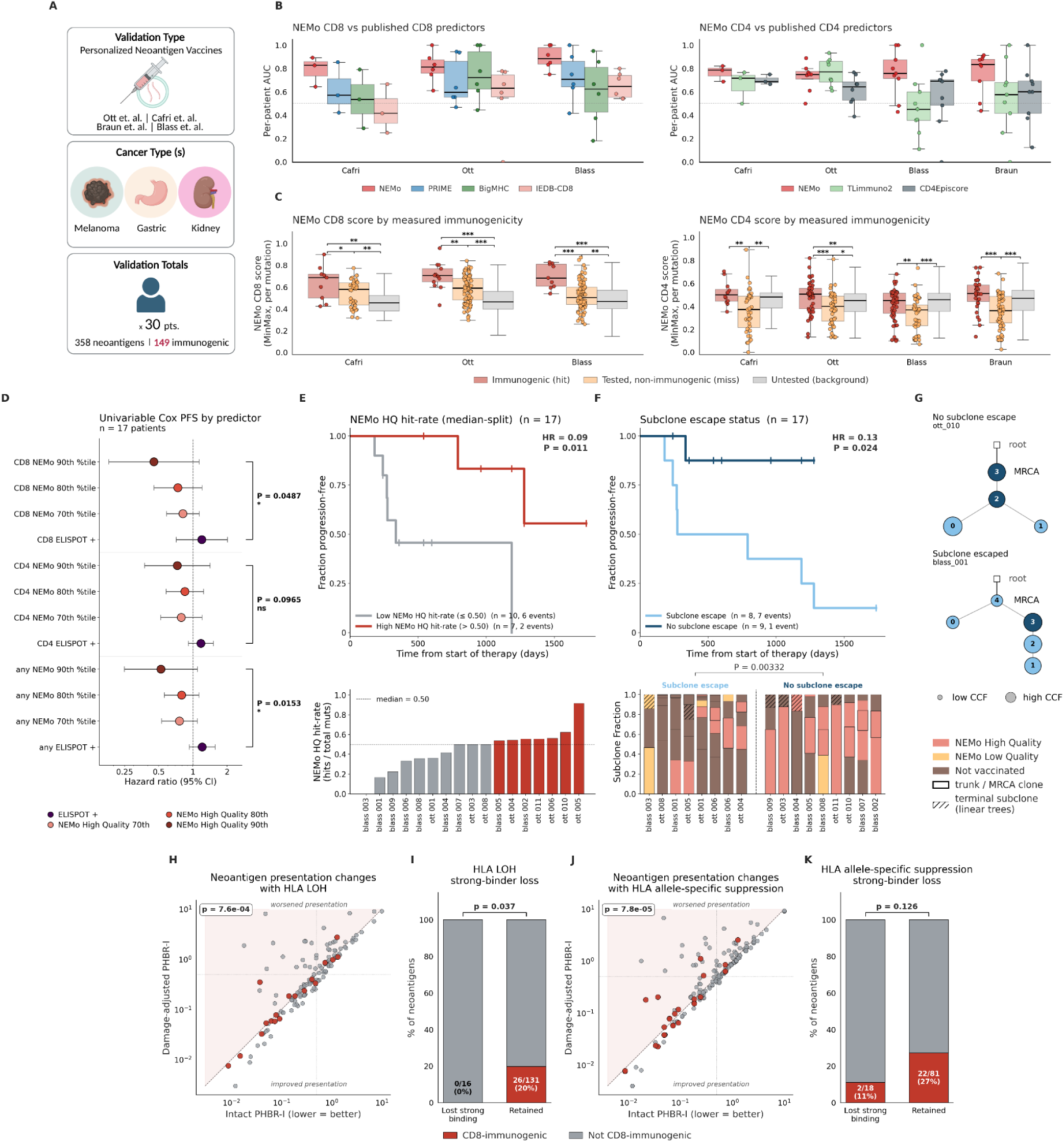
NEMo-predicted neoantigens track outcome in personalized cancer vaccine cohorts. Validation in personalized neoantigen-vaccine cohorts (Parkhurst/Cafri, Ott, Blass; Braun, CD4-only). **(A)** Schematic of cohorts and cancer types. **(B)** Per-patient AUC of NEMo versus comparator models (CD8: PRIME, BigMHC, IEDB-CD8; CD4: TLimmuno2, CD4Episcore). **(C)** Per-mutation NEMo score for assay-confirmed immunogenic peptides, screened-but-negative peptides, and unscreened background mutations, per cohort and per head. **(D)** Univariable Cox proportional hazards forest plot for progression-free survival (PFS, n = 17), displaying hazard ratios (HR, 95% CI) per additional covered hit across NEMo high-quality percentile thresholds versus ELISPOT positivity. **(E)** Kaplan–Meier curves of PFS stratified by median NEMo high-quality hit-rate split (higher curves indicate better outcome), with the lower panel illustrating per-patient high-quality hit fractions. **(F)** Kaplan–Meier analysis of PFS stratified by subclone escape status (“Subclone Escape” versus “No Subclone Escape”), alongside per-patient subclone vaccine coverage profiles. **(G)** Subclone-architecture schematics for a covered (ott_010) and an escaped (blass_001) tumor. **(H–K)** HLA LOH (H, I) and ASE (J, K) presentation-change scatter and strong-binder-loss bars. ASE, allele-specific expression.

Across all evaluable patients from the vaccine cohorts, NEMo produced a higher mean per-patient AUC than every comparator. These differences were significant for all CD8 comparators (paired two-sided t-test, P=1.58×10⁻³–0.0406) and for the recognition-aware CD4 comparators TLimmuno2 and CD4Episcore (P=0.00593 and 0.00463, respectively), but not for NetMHCIIpan or MixMHC2pred (P=0.360 and 0.287; Suppl. Fig. 8A,B). Descriptive mutation-pooled ROC analyses also placed NEMo first in both compartments (CD8 AUC=0.773; CD4 AUC=0.709; Suppl. Fig. 8C,D). Mean within-patient Spearman correlations indicated that NEMo generated complementary scoring profiles, with correlations ranging from ρ=0.17–0.66 for CD8 and ρ=0.09–0.68 for CD4 (Suppl. Fig. 8E,F). Per-mutation NEMo scores separated assay-confirmed hits from screened-but-negative mutations in every evaluable cohort and compartment (two-sided Mann–Whitney U, nominal P≤0.0495; Fig. 4C). Hit-versus-background separation was significant in all three CD8 cohorts and in Ott CD4 (P≤0.00383), but not in Blass CD4, Braun CD4, or Cafri CD4 (P=0.508, 0.0583, and 0.164, respectively; Suppl. Table 1). None of these studies prioritized CD4 responses in their vaccine designs, which may explain why many high-probability NEMo CD4 neoantigens remained unvaccinated against.

We then asked whether patients receiving vaccines with higher proportions of NEMo-prioritized neoantigens experienced better outcomes. This analysis was restricted to the 17 patients with screened immunogenicity data and progression-free survival (PFS) from the Ott and Blass cohorts; the Braun cohort was excluded because no progression events were observed. Patients with a NEMo high-quality (HQ) neoantigen fraction above the cohort median had longer PFS than those at or below the median (HR = 0.09, log-rank P = 0.011; Fig. 4E).

Univariable Cox models yielded protective hazard-ratio estimates for predicted-positive neoantigen counts at the 70th, 80th, and 90th percentiles, with the lowest estimates at the 90th percentile. In models containing both predicted-positive and empirical post-vaccine ELISpot-positive counts, their coefficients differed at the 90th percentile for joint NEMo and CD8, but not CD4 (two-sided Wald contrasts: P = 0.0153, 0.0487, and 0.0965, respectively; Fig. 4D). Extending from neoantigen quality to clonal architecture, patients classified as having no predicted subclone escape had longer PFS than those with predicted escape (HR = 0.13, log-rank P = 0.024; subclone-fraction comparison, Mann–Whitney U P = 0.00332; Fig. 4F), illustrated by covered (ott_010) and escaped (blass_001) phylogenies (Fig. 4G; definitions in Methods).

Applying the same patient-stratification procedure to comparator tools, 2 of 9 showed significant PFS associations based on predicted-positive fractions of neoantigens (PRIME and IEDB-CD8, log-rank P = 0.032 and 0.040, respectively; Suppl. Fig. 9). No comparator tool showed a significant PFS association using the corresponding subclone-escape classification (Suppl. Fig. 9).

Finally, we asked whether damage to presenting HLA alleles was associated with reduced detection of post-vaccination CD8 T-cell responses. After accounting for HLA LOH, no responses were detected against neoantigens predicted to lose strong-binding presentation (0/16), compared with responses against 26/131 (20%) neoantigens predicted to retain presentation (P = 0.037; presentation-change P = 7.6 × 10⁻⁴; Fig. 4H,I). Accounting for allele-specific downregulation revealed a similar, but nonsignificant, association: responses were detected against 2/18 neoantigens predicted to lose strong-binding presentation versus 28/81 predicted to retain it (P = 0.126; presentation-change P = 7.8 × 10⁻⁵; Fig. 4J,K). These findings associate compromised HLA presentation with a lower frequency of detectable CD8 responses, but do not establish whether the association reflects differences in T-cell priming, expansion, or persistence. These findings support that disruption of presenting HLA alleles serves as an active mechanism of neoantigen-specific immune escape[30] of direct relevance to the effectiveness of personalized cancer vaccines.

### NEMo identifies reactive TIL targets, and HLA escape explains who benefits

We applied NEMo to the Lowery cohort of NCI metastatic epithelial cancers treated with adoptive TIL therapy (34 patients; >5,000 neoantigens; 55 reactive; Fig. 5A). This cohort is a heterogeneous, heavily pre-treated population spanning multiple epithelial histologies, and a subset of patients were refractory to prior checkpoint blockade[2]. Tumors that have already progressed through immune pressure are expected to have edited or escaped their most visible antigens, and mixed histology introduces variation in mutational burden and baseline prognosis[31]. Despite this clinical heterogeneity, NEMo recovered reactive neoantigens well above chance (CD8 recall@20 = 0.41, n = 21; CD4 recall@20 = 0.55, n = 22; Fig. 5B) and significantly distinguished reactive from non-reactive targets in both compartments (CD8 P = 2×10⁻⁶; CD4 P = 4×10⁻⁸; Fig. 5C).

**Figure 5.**
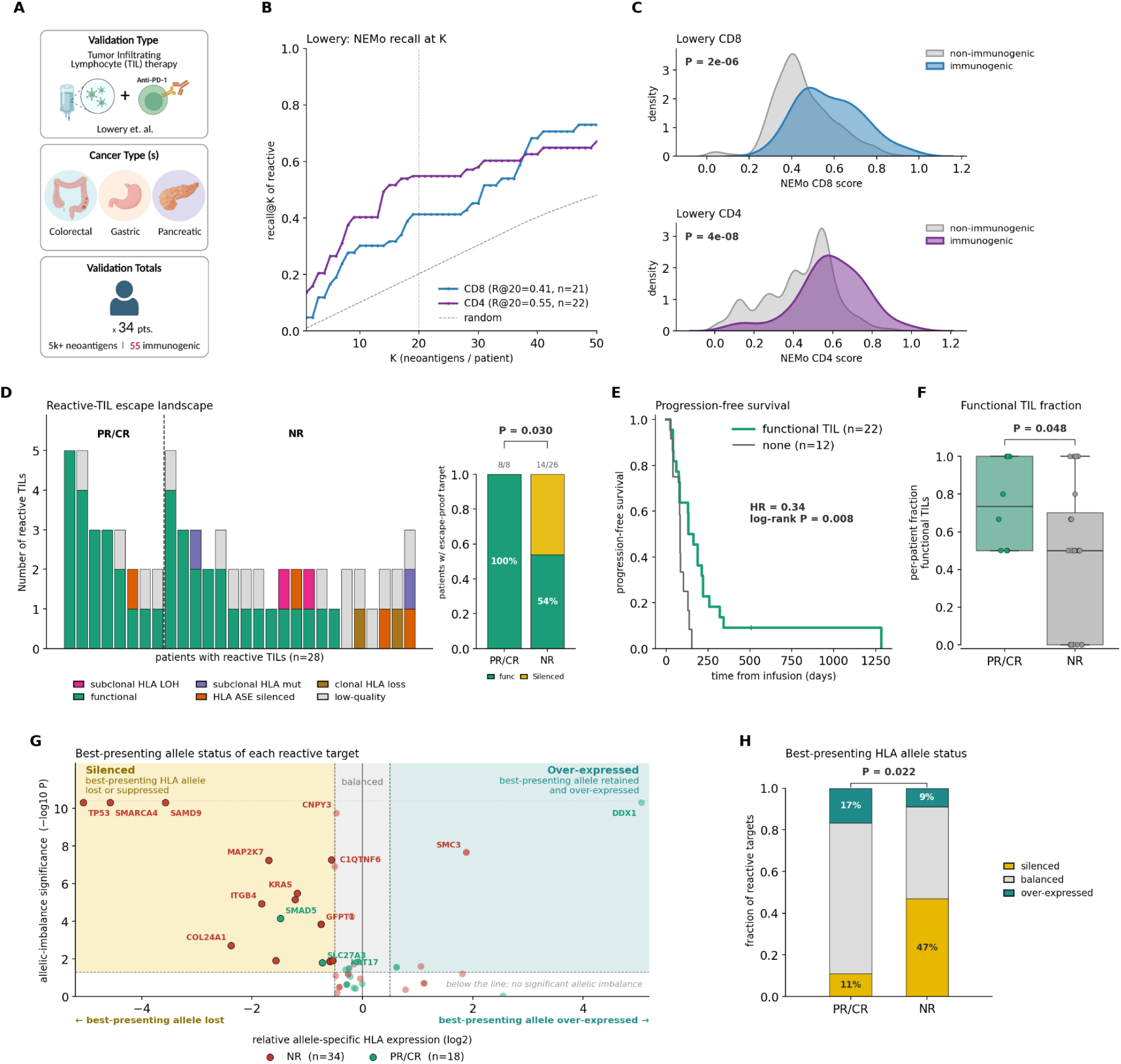
NEMo identifies reactive TIL targets, and HLA escape explains who benefits. Validation in the Lowery cohort of NCI metastatic-epithelial TIL therapy. **(A)** Schematic of cohort and cancer types. **(B)** Per-patient recall-at-K of reactive neoantigens for the CD8 and CD4 heads versus a random baseline. **(C)** NEMo-score separation of reactive versus non-reactive targets in the CD8 and CD4 compartments. **(D)** Per-patient reactive-TIL escape landscape, stratified by response (PR/CR vs NR) and colored by presenting-allele status (functional / HLA ASE-silenced / subclonal HLA LOH / subclonal HLA mutation / clonal HLA loss / low-quality). **(E)** Kaplan–Meier analysis of progression-free survival comparing patients with at least one escape-proof functional reactive TIL to those with none **(F)** Per-patient functional-TIL fraction by response. **(G)** Per reactive TIL, the expression imbalance of its presenting HLA allele (x-axis, log2 ratio of presenting to non-presenting allele; negative values indicate the presenting allele is under-expressed) against the significance of that imbalance (y-axis), colored by response. **(H)** Distribution of presenting HLA allele statuses (silenced, balanced, or over-expressed) for reactive targets in PR/CR versus NR tumors. PR, partial response; CR, complete response; NR, non-response.

In the original Lowery et al. study[2], TIL reactivity alone did not stratify patient survival. Significant associations with response and progression-free survival emerged only when we asked whether a patient’s reactive TILs were directed at targets the tumor could still present. Classifying every reactive TIL by its immune escape mechanism revealed that all responding patients (PR/CR; 8/8 100%) possessed at least one escape-proof functional reactive TIL, compared to only 54% (14/26) of non-responders (Fisher P = 0.030, Fig. 5D). Patients with at least one functional target achieved significantly longer PFS (HR = 0.34, log-rank P = 0.008; Fig. 5E), and the per-patient fraction of functional reactive TILs was higher in responders (P = 0.048, Mann-Whitney U; Fig. 5F).

Mapping each reactive TIL onto the expression imbalance of its presenting allele (Fig. 5G) showed HLA silencing (HLA lost or suppressed, Methods) concentrated in non-responders (47% loss) relative to responders (11% loss; χ² P = 0.022; Fig. 5H). TIL-reactive neoantigens were associated with a range of allele-specific HLA states, including loss or suppression of their predicted class-I presenting allele (Fig. 5G). Thus, experimentally detected T-cell recognition can coexist with alterations that may compromise tumor antigen presentation. This observation supports considering tumor presentation alongside neoantigen recognition when evaluating therapeutic targets[32]. However, these cross-sectional data do not establish whether HLA alterations preceded or followed immune recognition, or whether they directly limited treatment efficacy.

### Neoantigen editing is ICB-dependent: a ctDNA test of the training premise

While the preceding analyses establish that NEMo identifies clinically reactive targets, they do not directly establish whether the longitudinal training signal reflects immune selection or broader treatment effects. The SU2C-SARC032 (NCT03092323) trial and corresponding study[33] (hereafter termed the Subramanian cohort) provide a treatment-controlled setting to examine this distinction: patients received radiotherapy and surgery with or without perioperative pembrolizumab and underwent plasma sampling before treatment, after radiotherapy/before surgery, and at 3 and 12 months after surgery (Fig. 6A,B). Patients were identified using the clinical metadata and restricted to those with the sequencing inputs and mutation measurements required for this secondary analysis. Forty patients retained eligible mutations after filtering—15 receiving ICB and 25 controls. We tracked up to three prioritized mutations per patient (the dp3 set), alongside the remaining eligible nonsynonymous mutations and synonymous controls (Methods). Selecting up to three mutations focused tracking on the highest-priority candidates while applying a consistent selection rule across treatment arms. This choice was consistent with the limited breadth of experimentally detected neoantigen recognition in Parkhurst et al., which identified 0–3 TIL-recognized neoantigens in 92% of patients (69/75).

**Figure 6.**
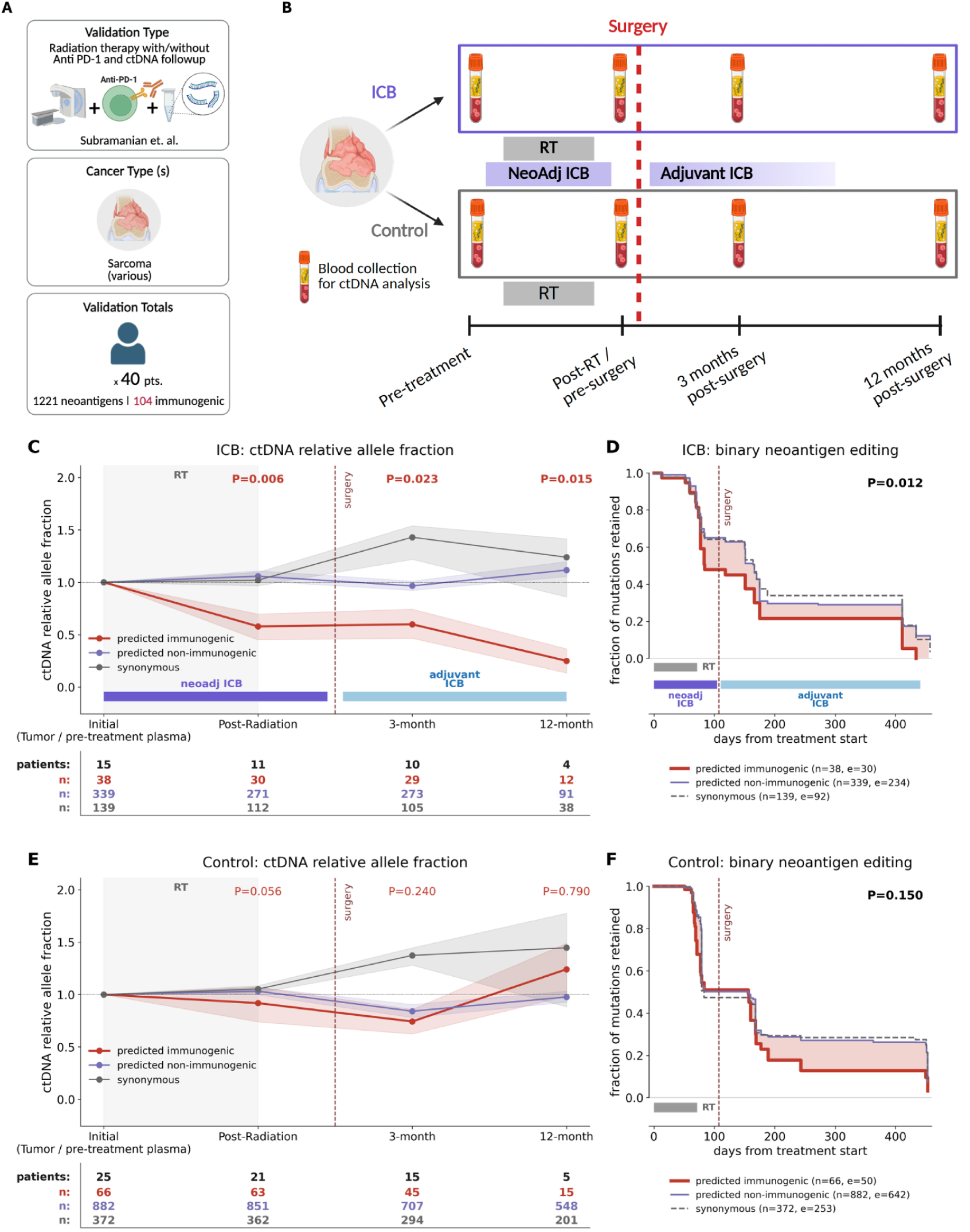
NEMo-prioritized mutations show relative depletion in the ICB-treated arm of the Subramanian ctDNA cohort. **(A)** Cohort schematic. **(B)** Trial design and plasma sampling timeline. Both arms received radiotherapy and surgery; the experimental arm additionally received neoadjuvant and adjuvant pembrolizumab. **(C)** Experimental ICB arm: mean patient-level relative allele-fraction fold change for NEMo-predicted immunogenic (dp3), predicted non-immunogenic, and synonymous mutations. Bands show pointwise 95% bootstrap confidence intervals. **(D)** Experimental ICB arm: Kaplan–Meier estimates of mutation retention, defined by remaining above the relative-depletion threshold (Methods). Shading highlights separation between immunogenic and non-immunogenic curves. **(E, F)** Corresponding allele-fraction and retention analyses in the radiotherapy-only control arm. P values compare predicted immunogenic and non-immunogenic mutations (Methods). ICB, immune checkpoint blockade; RT, radiotherapy; dp3, up to three NEMo-prioritized mutations per patient.

In the ICB-treated arm, circulating allele fractions of NEMo-predicted immunogenic mutations declined progressively relative to both predicted non-immunogenic and synonymous mutations, dropping to ∼27% of baseline values by 12 months (post-radiation P = 0.006; 3-month P = 0.023; 12-month P = 0.015; Fig. 6C), while non-immunogenic and synonymous fractions remained stable near 1.0. These per-timepoint tests are patient-level, as the number of ICB patients also declines across timepoints (15, 11, 10, 4). In addition to decreasing in relative abundance, predicted-immunogenic mutations were also eliminated significantly faster over time (P = 0.012; Fig. 6D; Methods). In the radiotherapy-only control arm, marginal or no such separation occurred at any timepoint (post-radiation P = 0.056; 3-month P = 0.240; 12-month P = 0.790; Fig. 6E), and elimination kinetics were not significant across mutation classes (P = 0.150; Fig. 6F), though non-significant separation occurred between predicted immunogenic vs. non-immunogenic neoantigens post surgery. Across comparator models, NEMo was the only tool to show significant preferential decline of prioritized mutations at all three follow-up timepoints and in the mutation-retention analysis in the ICB arm (Suppl. Fig. 10).

When evaluated in an independent clinical cohort and read out in plasma rather than tissue, NEMo separates immunogenic from non-immunogenic mutations only when checkpoint blockade is present. This supports the interpretation that the editing signal on which NEMo is trained reflects immune-mediated elimination.

## Discussion

Neoantigen prioritization has advanced from peptide–MHC binding prediction to models that learn molecular correlates of T-cell recognition, and these methods perform well against the assay-based labels on which they are trained. However, their common limitation is that such labels capture only the initial steps of a longer process: presentation, or reactivity measured in a reductionist system. We took a complementary approach by evaluating which neoantigens the immune system functionally eliminates from tumors in patients, training directly on this evolutionary signal. Despite a noisy training label and a modest internal cross-validated AUC, NEMo successfully recovered assay-confirmed immunogenic targets across spontaneous screening, personalized vaccine, adoptive TIL, and serial ctDNA cohorts in which it was never trained. Furthermore, NEMo outperformed comparator models that explicitly incorporate recognition features, and its predicted targets stratified progression-free survival more effectively than assay-confirmed reactivity.

The comparison with ELISPOT warrants careful consideration. ELISPOT detects IFN-γ secretion following ex-vivo peptide restimulation and is sensitive to experimental parameters including input cell number, in-vitro expansion protocol, peptide concentrations, and the threshold used to call a response. The assay measures whether a T-cell response is detectable under assay conditions, not whether that response is functional against tumor in vivo, nor does it account for whether the tumor retains the presenting HLA allele. Multimer-based enumeration, cytotoxicity assays, and TCR-repertoire tracking address parts of this limitation. Our finding that NEMo-predicted target coverage stratified survival better than ELISPOT positivity is therefore consistent with both the advantage of predicting targets that remain presentable by the tumor and with known limitations of the assay used as comparator, and we do not attempt to separate these contributions here.

A consistent architecture of immune evasion emerges from several independent directions. Oncogenic drivers are spared from editing and truncal mutations are protected relative to subclonal ones; neoantigens restricted by lost or down-regulated HLA alleles undergo less editing; experimentally confirmed immunogenic peptides in the screening cohorts are preferentially restricted by HLA alleles that are selectively silenced in the tumor; and in the TIL cohort HLA loss concentrates in non-responders. Together, these observations indicate that tumors reaching clinical detection have already resolved their most immunogenic mutations — either by never presenting them, or by immune escape — leaving immunoditing to act primarily on the non-driver, subclonal, still-presented periphery.

HLA-associated features showed contrasting associations with measured immunogenicity across validation settings. In Wells, validated neoantigens were enriched among peptides assigned to relatively lower-expressed HLA alleles (Fig. 3H). In the vaccine cohorts, CD8 T-cell responses were less frequently detected against mutations predicted to lose strong HLA presentation because of HLA loss of heterozygosity, with a similar but nonsignificant trend for allele-specific suppression (Fig. 4I,K). These observations are consistent with a context-dependent relationship between immune recognition and tumor escape, although the analyses measure different aspects of HLA-associated presentation.

These associations may reflect differences in how the studies identify immunogenic neoantigens. In the screening cohort, lower expression of the presenting HLA allele could reflect selection favoring tumor cells that evade an existing neoantigen-specific response. In the vaccine cohorts, responses were less frequently detected against targets predicted to lose presentation through HLA loss. One possible explanation is that reduced presentation by the tumor limits the stimulation or maintenance of vaccine-induced T cells. However, peptide-restimulation assays do not directly measure recognition of tumor cells, and nonmalignant antigen-presenting cells can support vaccine priming despite tumor-specific HLA loss. The present data therefore establish an association between predicted tumor presentation and detectable T-cell responses, but do not resolve its mechanism or temporal direction. Longitudinal measurements of antigen-specific T cells and direct recognition assays using autologous tumor cells would help distinguish these possibilities.

Although mutations in oncogenes and truncal mutations showed lower IVE-positive rates in the training cohort, neither group was uniformly spared from editing, and reactive TILs against TP53, KRAS, and PIK3CA demonstrate that driver-derived neoantigens can be recognized. Adoptive transfer delivers effector numbers far exceeding endogenous responses and may surface reactivity against weakly presented targets; alternatively, driver-directed responses may be preferentially class II–restricted and therefore outside the class-I editing signal. Distinguishing these possibilities would clarify whether driver mutations are permanently unavailable as targets or merely inaccessible under endogenous immune pressure, a question with direct bearing on shared-neoantigen therapeutic strategies.

These findings carry direct implications for personalized target prioritization. NEMo-predicted hits stratify survival where ELISPOT positivity and other computational tools do not, and in the TIL setting survival depends on escape-proof targets rather than merely reactive ones. Computational selection pipelines for vaccines and cell therapies should therefore explicitly weight presentation robustness — confirming the tumor retains specific alleles required to display the target—alongside predicted immunogenicity.

Several limitations should be noted. The in-vivo editing label is indirect, non-independent across mutations sharing a subclone, and subject to drift, sampling, and reconstruction error; internal cross-validated performance is measured against these same noisy labels and should not be read as an estimate of immunogenicity-prediction accuracy. Several validation cohorts are small (n = 17 for vaccine survival; n = 5 and n = 3 for the Wells analyses; n = 34 for TIL). Furthermore, the training cohort comprises ICB-treated tumors selected for evidence of immune activity, which may influence the observed patterns of editing across mutation classes. Whether the lower editing rates observed for oncogene and truncal mutations extend to primary-resistant or immunologically cold tumors remains unresolved. Comparisons across these settings could help distinguish intrinsic differences in immunogenicity from the effects of prior immune selection and current immune activity.

## Methods

### Training cohort and patient selection

The training set comprised 215 paired-timepoint patients on ICB with tumor whole-exome sequencing at two or more timepoints (pre- and post-treatment), and tumor RNA sequencing where available. Ninety-three patients (39,952 neoantigens) were classified as Included and retained for downstream model training following a curation cascade requiring evidence of active immune engagement — based on immunoediting, T-cell fraction, and CD274 (PD-L1) expression and their change between timepoints — together with quality control (QC) on age, tumor purity, and mutation burden. The remaining 122 patients were categorized as Excluded (n=47, failing active immune response criteria), QC failures (n=39, failing sequencing quality or clinical parameter thresholds), and Missing data (n=36, lacking sufficient data for analysis).

### Sequencing, variant calling, and neoantigen features

To ensure cross-study comparability, all cohorts were uniformly processed from raw sequencing reads using a unique pipeline. Tumor exomes—alongside matched normal and tumor RNA where available—were aligned to GRCh38, and somatic variants were called using nf-core/sarek (v3.4.2) with Mutect2 and annotated with VEP against Ensembl v111. Patient HLA class I and class II alleles were typed from exome data at four-digit resolution using HLA-HD (v1.7.0), establishing a uniform allele profile applied across NEMo and all comparator models. Per-mutation neoantigen features were extracted using NeoPrecis[25] across candidate mutant and wild-type 8–11mers (class I) and 15mers (class II) giving: PHBR-I and PHBR-II to quantify presentation; agretopicity to evaluate mutant versus wild-type binding affinity; foreignness to capture alignment to known immunogenic epitopes; cross-reactivity distance to capture TCR-contact divergence; robustness to quantify multi-allele coverage; and RNA expression metrics. NEMo then integrates all this information alongside additional features to model the full in-vivo editing cascade.

### Subclonal reconstruction, CCF, and the editing (IVE) label

Subclonal structure and per-mutation cancer cell fraction (CCF) were reconstructed with PhyClone across all available timepoints. For each (patient, mutation) a closed-form ordinary-least-squares slope of CCF against biopsy index was fit over all timepoints. A mutation was labeled IVE-positive (High_to_Low) when it satisfied all three of: slope ≤ −0.075, dynamic range (max − min CCF) ≥ 0.25, and initial CCF ≥ 0.10. The mirror class (Low_to_High) required slope ≥ +0.075, dynamic range ≥ 0.25, and initial CCF ≤ 0.50; remaining mutations were labeled Other and collapsed with Low_to_High for training. Mutations with both initial CCF < 0.10 and max CCF < 0.10 across all timepoints were dropped as deconvolution noise. Config: SLOPE_NEG_THRESH = −0.075, SLOPE_POS_THRESH = 0.075, DYN_RANGE_THRESH = 0.25, MIN_INITIAL_HIGH = 0.10, MAX_INITIAL_LOW = 0.50, DROP_LOW_LOW = True, LOW_LOW_MAX_CCF = 0.10. Editing rate denotes the proportion of mutations within a stratum meeting the IVE-positive definition. Subclone depth denotes the number of edges separating a mutation’s subclone from the trunk (MRCA) in the PhyClone tree, and is independent of cancer cell fraction.

### NEMo model

NEMo comprises two machine learning classifiers, one per T-cell compartment—a CD8 head over MHC class-I peptides and a CD4 head over class-II peptides. The two heads are trained and evaluated independently because class-I and class-II peptides differ in length and binding rules; cross-compartment comparisons are therefore made over the same genomic region rather than the same peptide. Models were trained on the editing label and evaluated by 10-fold cross-validation. A full description of NEMo architecture will be made available upon formal publication.

### Immunogenicity ground truth by cohort

Screening cohorts (Parkhurst, Wells) measure pre-existing reactivity: candidate neoantigens are synthesized and assessed for recognition by patient-derived T cells, in the Wells cohort using barcoded peptide–MHC multimers. Vaccine cohorts (Parkhurst/Cafri, Ott, Blass, Braun) measure vaccine-induced reactivity by post-immunization IFN-γ ELISPOT on PBMCs, with intracellular cytokine staining used in some studies for CD4/CD8 attribution. TIL cohort (Lowery) reactivity was determined by TMG/ELISPOT screening of expanded TIL products. Because assay format and the meaning of a positive label differ across settings, per-cohort performance is reported separately throughout.

## Benchmarking tools

### Immunogenicity prediction

NEMo was benchmarked against PRIME v2.1, BigMHC-IM, and IEDB class-I immunogenicity release v3.0 for CD8 prediction, and TLimmuno2 and CD4Episcore for CD4 prediction.

PRIME used MixMHCpred v3.0 with mutant peptide sequences and patient-specific HLA-I alleles. BigMHC was run in immunogenicity mode using mutant peptide–HLA-I pairs. IEDB class-I immunogenicity received peptide sequences with default settings and without allele-specific conditioning. TLimmuno2 received mutant 15-mers paired with supported patient HLA-II alleles. CD4Episcore was accessed through the IEDB Next-Generation API using the cd4episcore method and 15-mer inputs. CD4Episcore provides population-level predictions rather than patient-specific HLA-restricted predictions.

### Binding-affinity and presentation prediction

Additional comparators comprised NetMHCpan v4.2, NetMHCIIpan v4.3e, MHCflurry v2.0.6 in presentation mode, and MixMHC2pred v2.0. Allele-dependent tools received mutant peptides and supported patient-specific HLA alleles. NetMHCpan and NetMHCIIpan were evaluated using BA percentile ranks over EL ranks where validation datasets overlapped with training.

### Mutation-level aggregation

Each comparator independently selected its most favorable prediction among the evaluated peptides—and patient HLA alleles where applicable—mapping to each mutation: the highest immunogenicity or presentation score, or the lowest percentile rank. Scores were oriented so that higher values indicated greater predicted immunogenicity or presentation for performance calculations.

### Evaluation metrics and statistics

Recall@K is the primary metric throughout. For a given head, it is the fraction of a patient’s confirmed immunogenic/reactive neoantigens falling within the K top-scoring candidates, macro-averaged across patients so that each patient contributes equally regardless of mutational burden, against a random-expectation baseline of min(1, K/n) for a patient with n scored candidates. Specifically, we report recall@20 as the primary operating point because current personalized cancer vaccines carry on the order of twenty neoantigens per patient[1], [14]. Full recall-at-K curves are nonetheless reported so performance is not read from a single cutoff, with K = 20 marked. Per-patient AUC was computed as the mean over patients contributing at least one positive and one negative peptide to the head-matched evaluation pool (class-I peptides for the CD8 head, class-II for CD4). Tests were Mann–Whitney U, log-rank and Cox proportional hazards (lifelines), Fisher exact, chi-square and chi-square-trend, and Wilcoxon signed-rank for paired timepoints; p-values and hazard ratios are reported in-figure.

### Vaccine subclonal coverage and end-lineage coverage

For each patient we compute the fraction of the tumor “covered” by the vaccine, where a subclone counts as covered if it carries at least one NEMo high-quality predicted neoantigen. The KM median split (Fig. 4E) uses hit_rate_any — the fraction of a patient’s mutations that are NEMo high-quality hits — split at the cohort median (∼0.50). The subclone-escape analysis (Fig. 4F) additionally uses a prevalence-weighted coverage (covered_prev_frac = Σ clonal_prev over covered clones), so covering a widespread subclone counts more than covering a rare one.

End-lineage coverage asks where coverage falls: whether predicted targets reach the terminal (leaf) subclones at the tips of the evolutionary tree rather than only the ancestral trunk. Tree structure is taken from the PhyClone Newick tree (trunk/MRCA = the single numeric child of the root; leaf = terminal clone; linear tree = no branching). End-lineage coverage is operationalized in the escape rule, which classifies each patient as escape− (contained) or escape+ (escaped): (i) a linear tree whose terminal leaf is hit is classified as escape−; (ii) when the trunk is hit, each branch off the trunk is compared for uncovered versus covered prevalence, with the trunk given ancestral credit scaled by √(branch size), and any branch with more uncovered than covered prevalence makes the patient escape+; (iii) when the trunk is missed, a branched tree is classified as escape+, whereas a linear tree is decided by summed covered versus uncovered prevalence along the chain.

### Subramanian et al. ctDNA analysis

#### Cohort and data source

We performed a secondary analysis of tumor, matched-normal, and plasma sequencing data from SU2C-SARC032 (NCT03092323), obtained from dbGaP (phs003921.v2.p1). Trial design, eligibility, specimen collection, personalized hybrid-capture panel design, and the original ctDNA assay are described by Subramanian et al.[33]. Briefly, patients received preoperative radiotherapy and surgery with or without perioperative pembrolizumab, with plasma sampling before treatment, after radiotherapy/before surgery, and at 3 and 12 months after surgery. Our analysis required pre-treatment tumor-derived candidates, a targeted pre-treatment tumor reference, and available NEMo predictions for nonsynonymous mutations. Patients with only post-radiotherapy tumor data were excluded. Targeted tumor depth ≥30 and VAF 0.05–0.90 were required, leaving 40 patients (15 ICB, 25 control). Follow-up availability varied because of missing draws, unavailable sequencing measurements, and insufficient coverage.

#### Secondary sequencing analysis

Candidate mutations were obtained from independently generated WES-derived NeoPrecis tables. Synonymous comparators were selected from VEP-annotated Mutect2 calls, excluding neoantigen loci and protein-altering variants, and subjected to the same targeted tumor depth and VAF thresholds. All retained mutations were SNVs. Targeted reads were aligned to hg38 using BWA-MEM and processed with GATK MarkDuplicates. Mutation VAFs were recalculated from alternate-base counts and filtered pysam pileup depth, using base quality ≥20 and excluding duplicate-flagged, secondary, unmapped, and QC-failed reads; nonsynonymous counting additionally excluded mapping-quality-zero reads. Plasma measurements required depth ≥100. Zero alternate reads at adequately covered loci yielded VAF = 0; missing or insufficient-depth measurements remained missing. Clinically ctDNA-negative samples were not included in the analysis.

#### NEMo prioritization and normalization

Up to three mutations per patient were selected using both NEMo prediction heads (dp3). Candidates with RNA TPM>0 formed the ranking pool when at least four were available; otherwise, all eligible mutations were ranked. Mutations at or above the respective 80^th^ percentiles for both prediction heads were prioritized, followed by the better rank across heads and then the sum of ranks. Remaining eligible nonsynonymous mutations formed the comparison group.

Relative allele fraction (relAF) was calculated using a shared patient/timepoint denominator: the mean VAF across all evaluable eligible nonsynonymous and synonymous mutations. The same normalization was applied to targeted pre-treatment tumor measurements. Patient-class mean relAF values were expressed as fold changes from the corresponding positive pre-treatment plasma mean, with normalized targeted-tumor fallback when the plasma reference was missing or nonpositive. Zero normalization denominators yielded missing values. Fold changes were averaged with equal patient weight.

#### Endpoints and statistics

Longitudinal analyses required a positive initial reference and at least one evaluable follow-up measurement. Pointwise bootstrap intervals used 1,000 resamples of whole patient trajectories and reported 95% confidence intervals. Per-timepoint comparisons used two-sided, unpaired Mann–Whitney U tests for lower dp3 fold changes.

Kaplan–Meier retention analyses defined an event as the first evaluable follow-up relAF strictly below 50% of the mutation’s positive initial plasma reference, with targeted-tumor fallback when required. Missing visits were skipped, mutations without events were censored at their last evaluable draw, and later reappearance did not reverse an event. Event times used recorded days from treatment start. Dp3 and remaining nonsynonymous mutations were compared using a one-sided log-rank test with 2,000 within-patient label permutations.

## Code Availability

All code will be available upon formal publication.

## Data Availability

All datasets used and corresponding accession numbers (if applicable) will be referenced upon formal publication.

## Acknowledgements

We are deeply grateful to the patients and their families whose participation made this research possible, and to the investigators and study teams who generated and shared the validation datasets used here, including the TESLA consortium (Wells et al.) and the studies by Parkhurst, Lowery, Cafri, Ott, Blass, and Braun and colleagues. This work was funded by Mark Foundation Emerging Leader Award #18-022-ELA, NCI grant R01CA269919, and support from NCI grant U24CA248138 to H. Carter. Computational resources were supported by infrastructure grant 2P41GM103504-11. We thank the Moding lab at Stanford for their close collaboration with the ctDNA analysis. We also thank Jenna Chen for her contributions to figure and schematic design. Full acknowledgements of the participating studies and contributing authors will accompany formal publication.

## Supplementary Figures

**Supplementary Figure 1.**
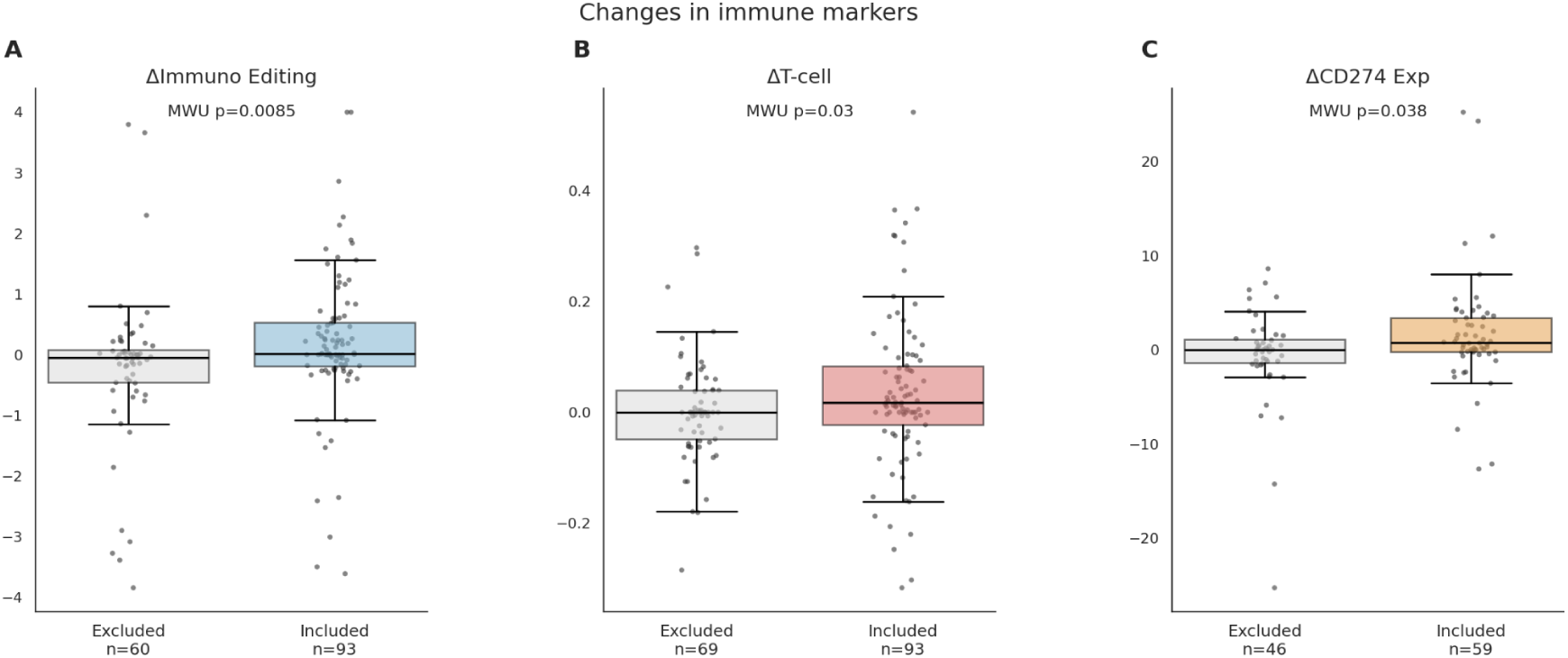
Changes in immunoediting (A), T-cell measurements (B), and CD274 expression (C) in included and excluded patients with available paired measurements. Immunoediting change is displayed as TP1 minus TP2; other changes are TP2 minus TP1. Groups are compared by Mann–Whitney U tests; sample sizes are indicated. Excluded patients comprise all other paired-cohort patients with the relevant measurements.

**Supplementary Figure 2.**
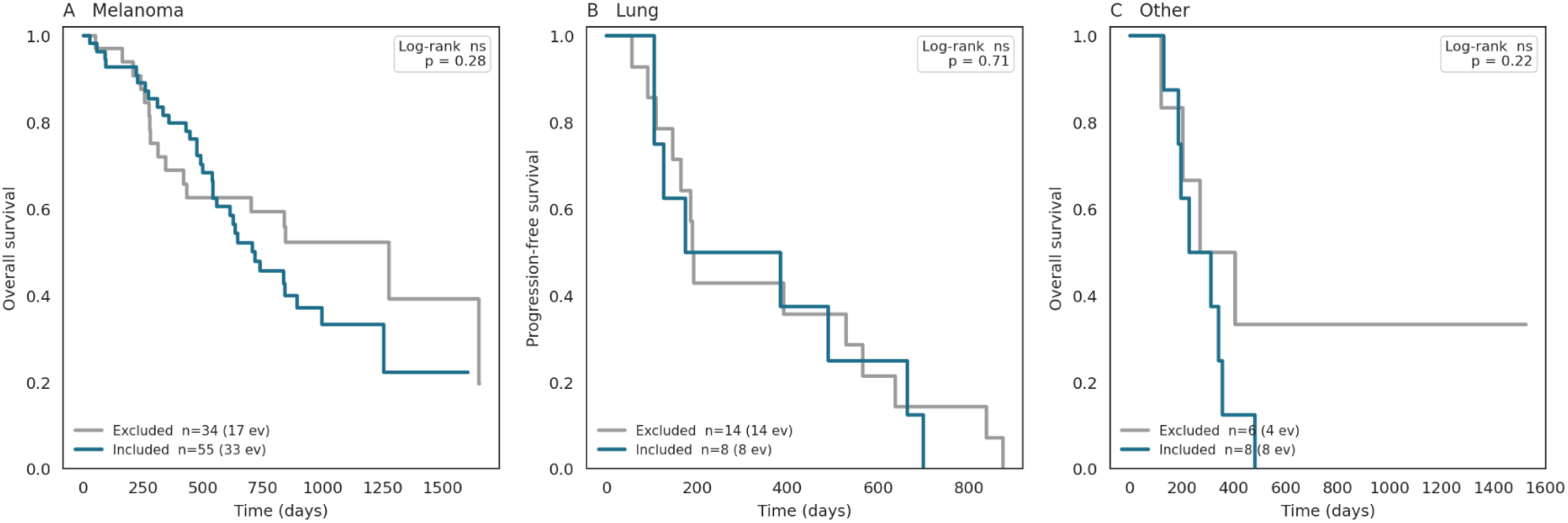
Kaplan–Meier survival estimates for included and excluded patients with melanoma (A), lung cancer (B), and other cancers (C). Overall survival is shown in A and C and progression-free survival in B; groups are compared by log-rank tests. Patient and event counts are indicated. Lung curves are an all-event sensitivity analysis in which every recorded follow-up time is treated as an event, without censoring.

**Supplementary Figure 3.**
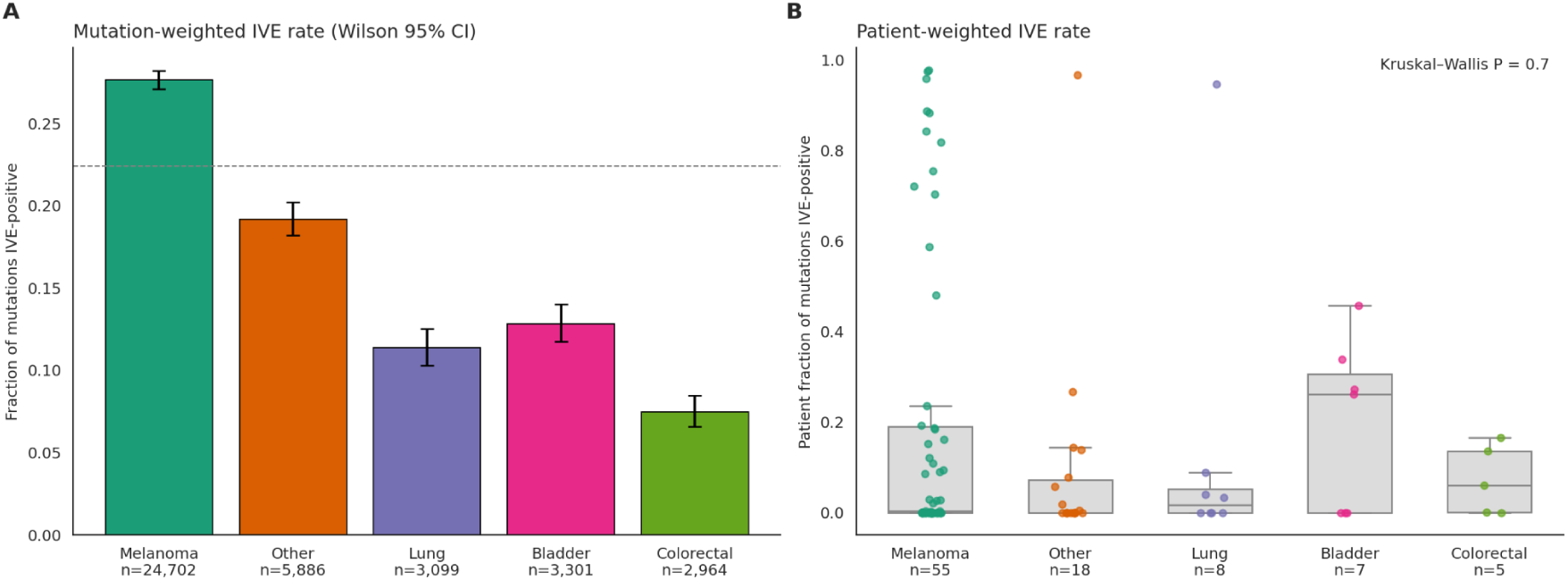
(A) Fraction of mutations positive for in vivo editing (IVE) by cancer type, with Wilson 95% confidence intervals; the dashed line indicates the overall mutation-level rate. (B) Per-patient IVE-positive fractions. Points represent patients; boxes show medians and interquartile ranges, with whiskers extending to observations within 1.5 times the interquartile range. Cancer types are compared by a Kruskal–Wallis test. Mutation counts (A) and patient counts (B) are indicated.

**Supplementary Figure 4.**
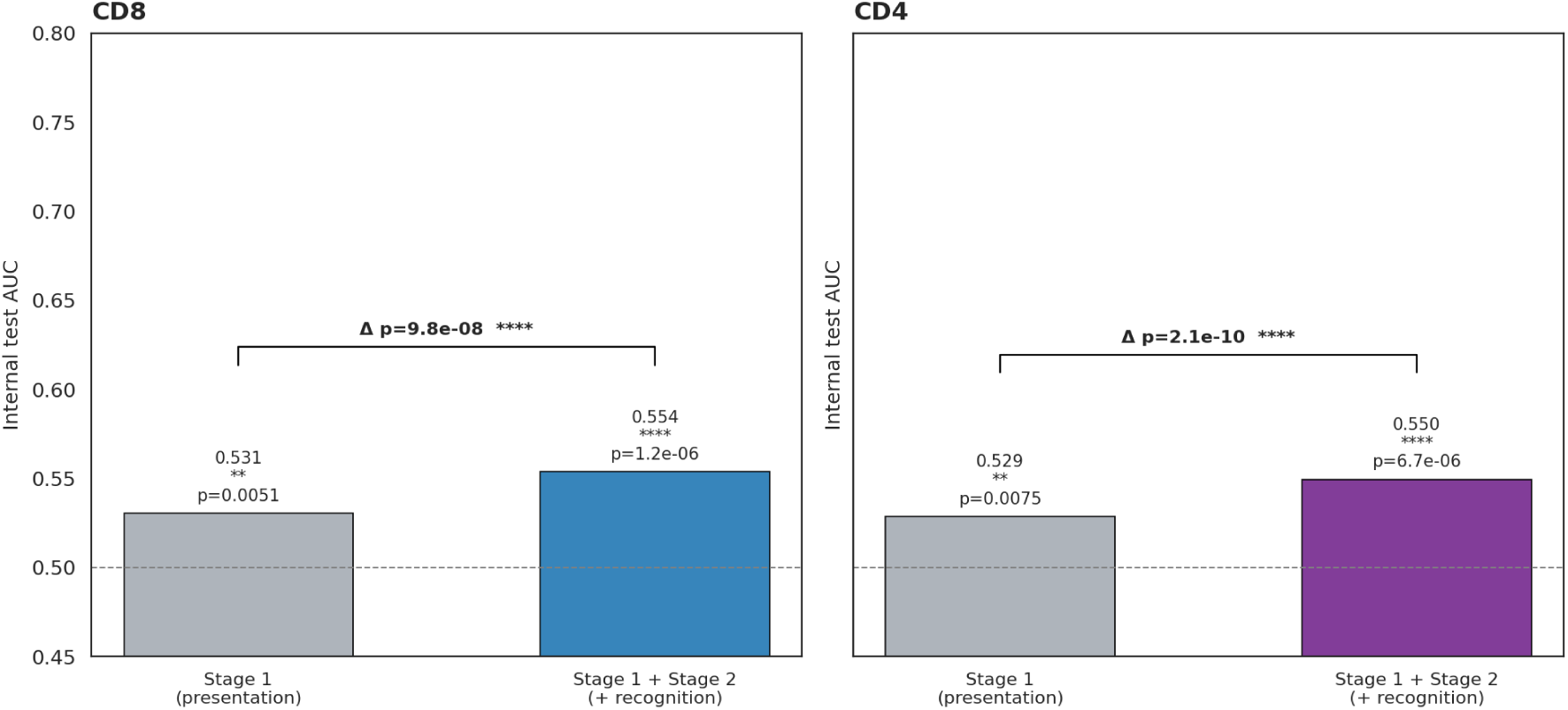
Held-out test-set AUC for the presentation-only stage and the full presentation-plus-recognition model, shown separately for CD8 and CD4. Paired DeLong tests compare stages on the same test observations; annotations above each bar test AUC against 0.5. Saved models from the pinned analysis run were used without retraining.

**Supplementary Figure 5.**
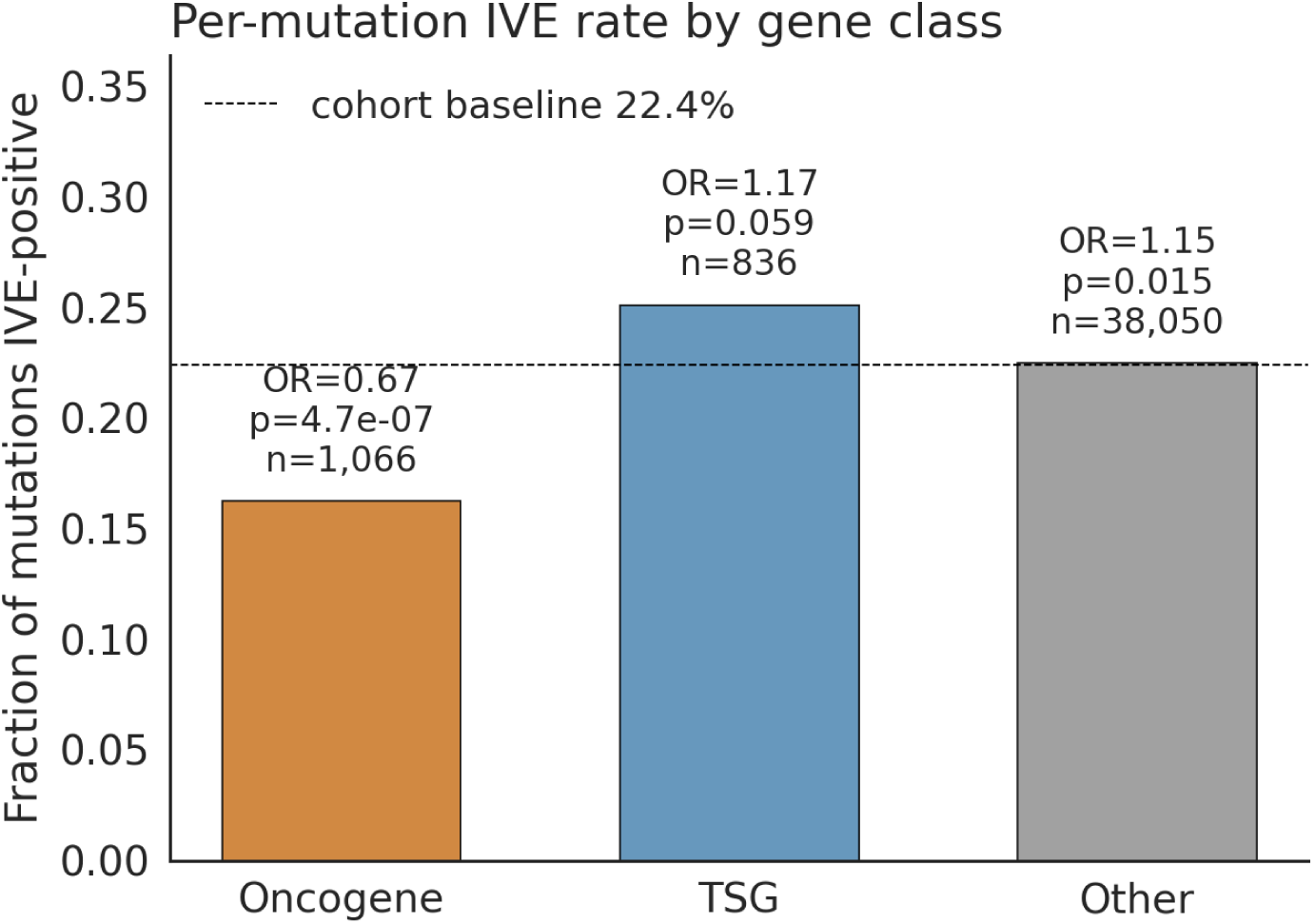
Mutation-level IVE-positive fractions in oncogenes, tumor-suppressor genes (TSG), and other genes. The dashed line indicates the cohort-wide rate. Odds ratios and two-sided Fisher exact P values compare each class with all remaining mutations; mutation counts are indicated. Comparisons are unadjusted.

**Supplementary Figure 6.**
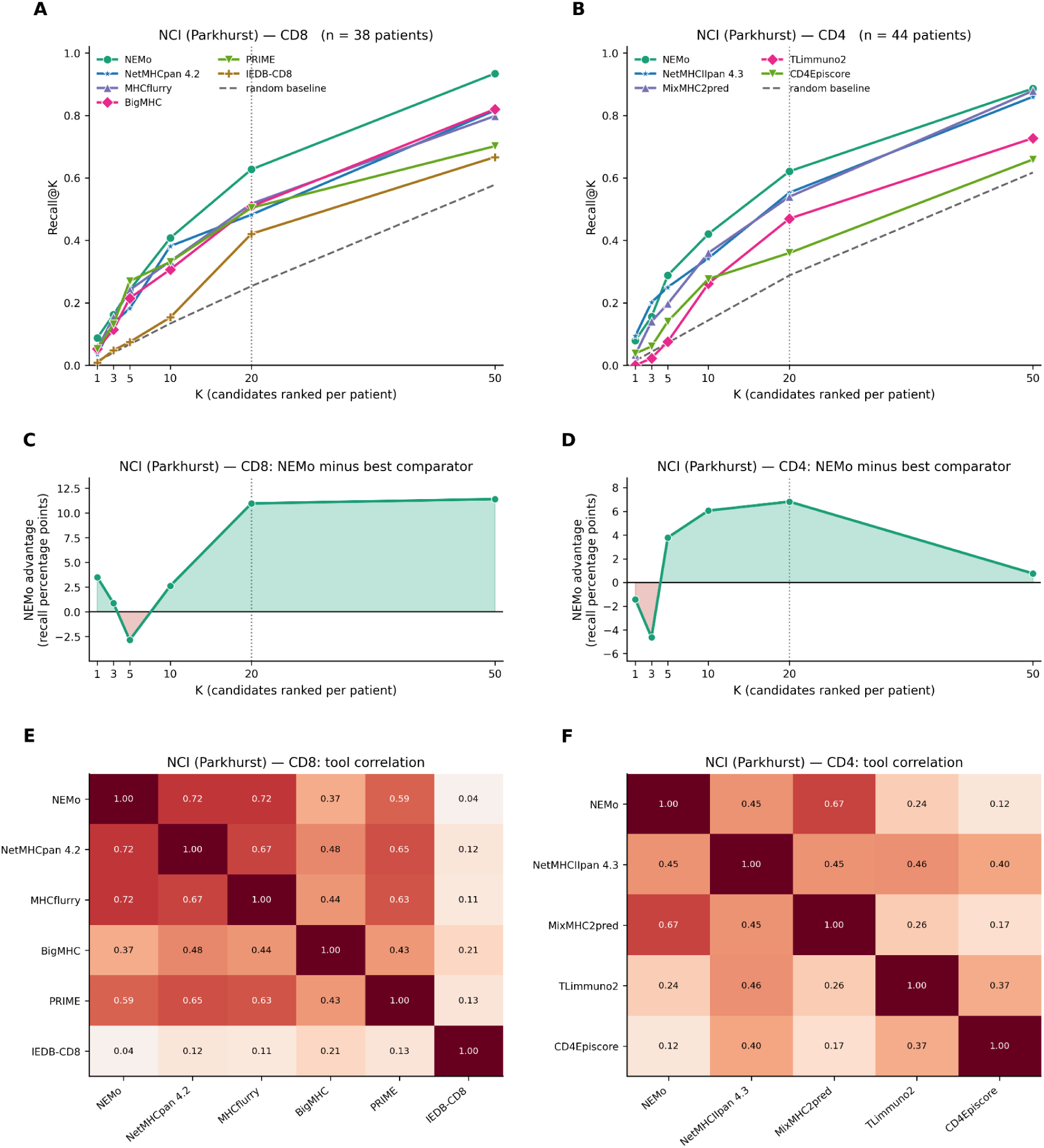
(A,B) Mean per-patient recall at K for CD8- and CD4-reactive mutations, respectively, within the 75-patient Parkhurst cohort. Recall includes patients with at least one retained reactive mutation in the relevant compartment (38 CD8; 44 CD4); dashed curves show random ranking. (C,D) NEMo recall minus the highest comparator recall at each K, expressed in percentage points. (E,F) Mean within-patient Spearman correlations between tool scores. Vertical references mark K = 20.

**Supplementary Figure 7.**
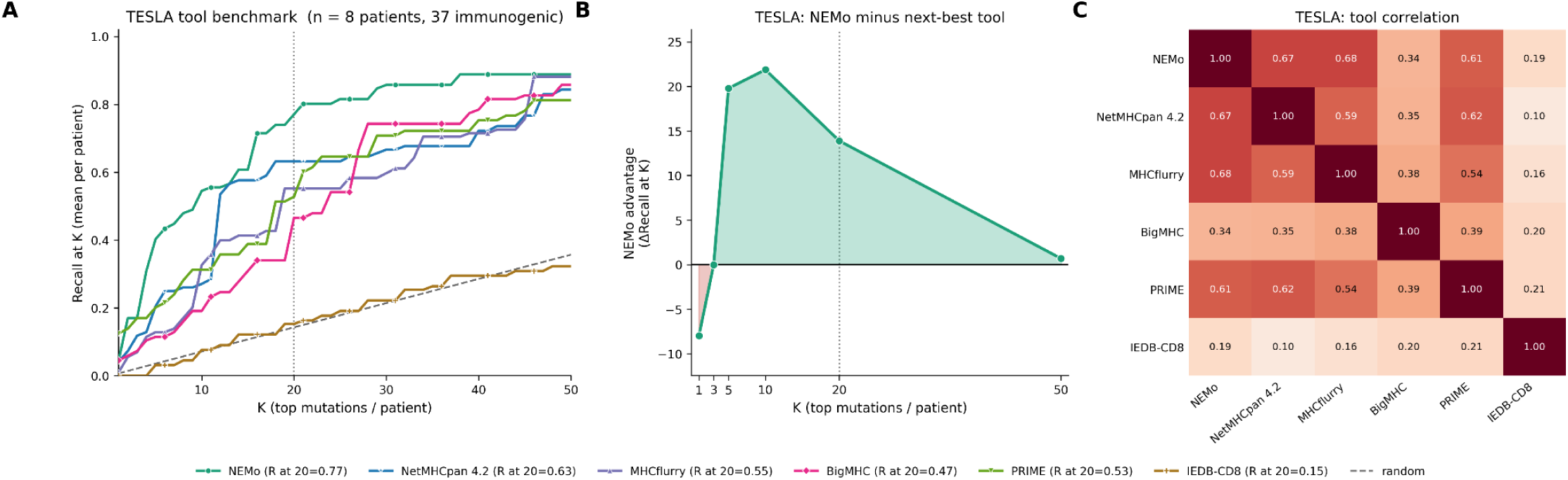
(A) Mean per-patient recall at K for NEMo and comparator tools across eight patients with 37 immunogenic mutations. The dashed curve indicates random ranking. (B) NEMo recall minus the highest comparator recall at each K, expressed in percentage points. (C) Mean within-patient Spearman correlations between tool scores. K denotes mutations ranked per patient; the vertical reference marks K = 20.

**Supplementary Figure 8.**
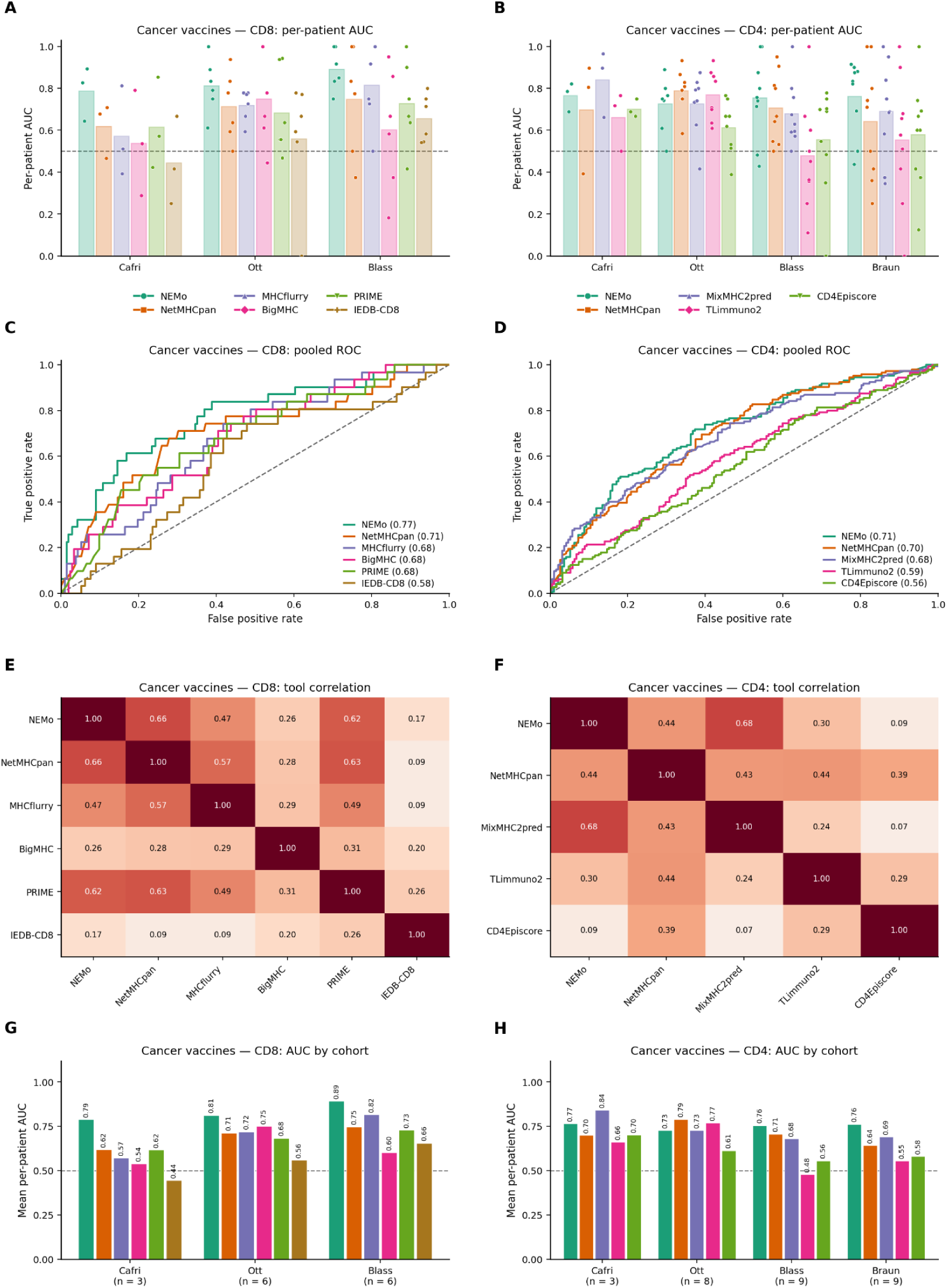
(A,B) Per-patient AUCs for CD8 and CD4 across cancer-vaccine cohorts; points represent patients and bars indicate means. (C,D) ROC curves pooling mutations across cohorts, with pooled AUCs indicated. (E,F) Mean within-patient Spearman correlations between tool scores. (G,H) Mean per-patient AUC by tool and cohort, with contributing patient counts. Pooled ROC analyses are descriptive and do not establish patient-level superiority over each comparator.

**Supplementary Figure 9.**
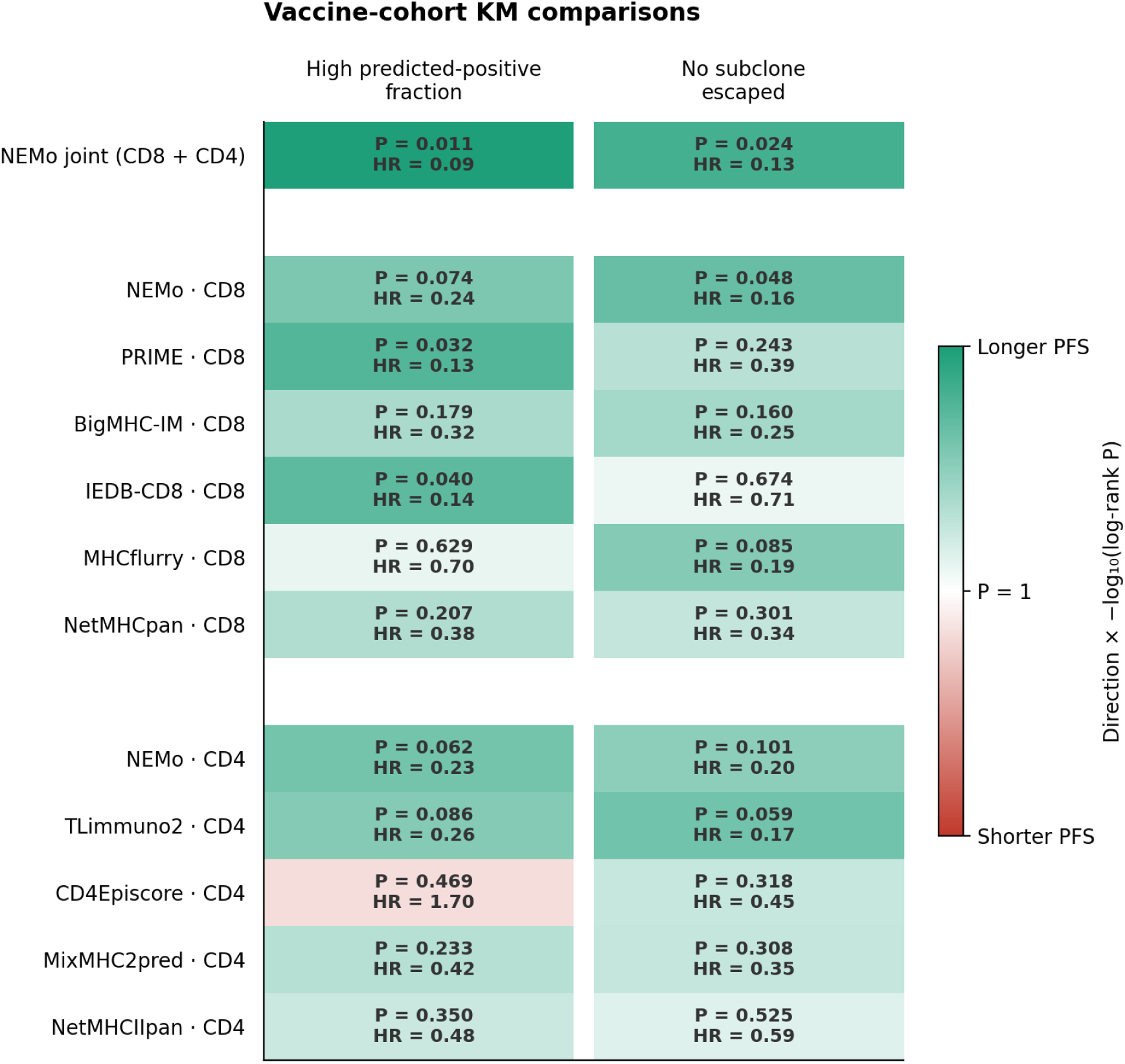
Heatmap of progression-free survival (PFS) comparisons in the Blass and Ott vaccine cohorts using the Figure 4 stratification procedures (17 patients, 8 progression events). Patients were stratified by the fraction of clone-mapped vaccine mutations classified as predicted-positive (above versus at or below the tool-specific median; left) or by predicted subclone escape (absent versus present; right). Positive calls used each tool’s oriented-score 70th percentile in the pooled vaccine-mutation reference set; joint NEMo calls met either head’s threshold. Subclone escape was assigned using the Figure 4 phylogenetic rule. Cells show two-sided log-rank P values and hazard ratios (HRs) from univariable Cox models. Green indicates lower hazard in the high-fraction or no-escape group; red indicates higher hazard. Color intensity represents negative log10 log-rank P. White gaps separate joint NEMo, CD8, and CD4 analyses. P values are nominal.

**Supplementary Figure 10.**
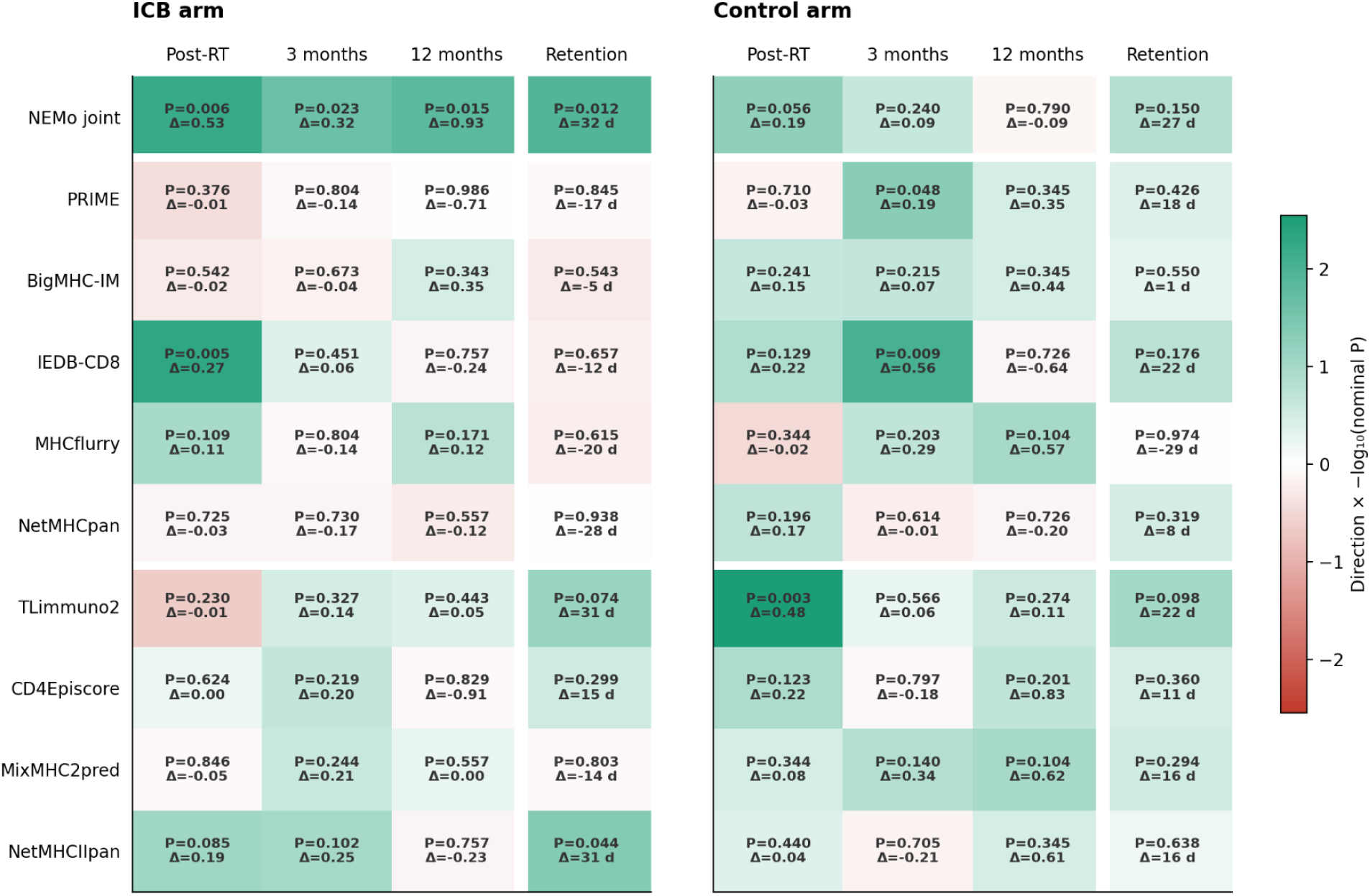
Comparison of tool-prioritized mutations under shared ctDNA normalization. Each tool prioritized up to three mutations per patient across 40 patients. Heatmaps summarize preferential decline of selected versus remaining nonsynonymous mutations in the ICB (left) and control (right) arms. VAFs were normalized to the patient-specific mean across all eligible nonsynonymous and synonymous mutations before tool-specific grouping. Follow-up comparisons used baseline-anchored, patient-level relative allele fractions (Mann–Whitney U tests). Mutation retention was defined by the first observed decline below 50% of normalized baseline and compared using 2,000 within-patient label permutations of a one-sided log-rank statistic. Δ denotes the median difference in relative allele fraction, or the difference in restricted mean retention time through day 365 (other minus selected); positive values indicate preferential decline of selected targets. Colors encode sign(Δ) × −log₁₀(P). Displayed P values are nominal.

## References

[1] L. A. Rojas et al., “Personalized RNA neoantigen vaccines stimulate T cells in pancreatic cancer,” Nature, vol. 618, no. 7963, pp. 144–150, Jun. 2023.

[2] F. J. Lowery et al., “Neoantigen-specific tumor-infiltrating lymphocytes in gastrointestinal cancers: a phase 2 trial,” Nat. Med., vol. 31, no. 6, pp. 1994–2003, Jun. 2025.

[3] R. Leidner et al., “Neoantigen T-cell receptor gene therapy in pancreatic cancer,” N. Engl. J. Med., vol. 386, no. 22, pp. 2112–2119, Jun. 2022.

[4] N. McGranahan and C. Swanton, “Neoantigen quality, not quantity,” Sci. Transl. Med., vol. 11, no. 506, p. eaax7918, Aug. 2019.

[5] M. Łuksza et al., “Neoantigen quality predicts immunoediting in survivors of pancreatic cancer,” Nature, vol. 606, no. 7913, pp. 389–395, Jun. 2022.

[6] Y. Wu et al., “The predictive value of tumor mutation burden on efficacy of immune checkpoint inhibitors in cancers: A systematic review and meta-analysis,” Front. Oncol., vol. 9, p. 1161, Nov. 2019.

[7] F. Huber et al., “A comprehensive proteogenomic pipeline for neoantigen discovery to advance personalized cancer immunotherapy,” Nat. Biotechnol., vol. 43, no. 8, pp. 1360–1372, Aug. 2025.

[8] L. Lybaert et al., “Challenges in neoantigen-directed therapeutics,” Cancer Cell, vol. 41, no. 1, pp. 15–40, Jan. 2023.

[9] N. Xie, G. Shen, W. Gao, Z. Huang, C. Huang, and L. Fu, “Neoantigens: promising targets for cancer therapy,” Signal Transduct. Target. Ther., vol. 8, no. 1, p. 9, Jan. 2023.

[10] M. R. Parkhurst et al., “Unique neoantigens arise from somatic mutations in patients with gastrointestinal cancers,” Cancer Discov., vol. 9, no. 8, pp. 1022–1035, Aug. 2019.

[11] D. K. Wells et al., “Key Parameters of Tumor Epitope Immunogenicity Revealed Through a Consortium Approach Improve Neoantigen Prediction,” Cell, vol. 183, no. 3, pp. 818–834.e13, Oct. 2020.

[12] P. A. Ott et al., “An immunogenic personal neoantigen vaccine for patients with melanoma,” Nature, vol. 547, no. 7662, pp. 217–221, Jul. 2017.

[13] E. Blass et al., “A multi-adjuvant personal neoantigen vaccine generates potent immunity in melanoma,” Cell, vol. 188, no. 19, pp. 5125–5141.e27, Sep. 2025.

[14] D. A. Braun et al., “A neoantigen vaccine generates antitumour immunity in renal cell carcinoma,” Nature, vol. 639, no. 8054, pp. 474–482, Mar. 2025.

[15] G. Cafri et al., “mRNA vaccine-induced neoantigen-specific T cell immunity in patients with gastrointestinal cancer,” J. Clin. Invest., vol. 130, no. 11, pp. 5976–5988, Nov. 2020.

[16] B. C. Creelan et al., “Tumor-infiltrating lymphocyte treatment for anti-PD-1-resistant metastatic lung cancer: a phase 1 trial,” Nat. Med., vol. 27, no. 8, pp. 1410–1418, Aug. 2021.

[17] J. B. Nilsson, J. Greenbaum, B. Peters, and M. Nielsen, “NetMHCpan-4.2: improved prediction of CD8+ epitopes by use of transfer learning and structural features,” Front. Immunol., vol. 16, no. 1616113, p. 1616113, Aug. 2025.

[18] J. B. Nilsson et al., “Accurate prediction of HLA class II antigen presentation across all loci using tailored data acquisition and refined machine learning,” Sci. Adv., vol. 9, no. 47, p. eadj6367, Nov. 2023.

[19] D. J. Schwartzentruber et al., “Gp100 peptide vaccine and interleukin-2 in patients with advanced melanoma,” N. Engl. J. Med., vol. 364, no. 22, pp. 2119–2127, Jun. 2011.

[20] J. Schmidt et al., “Prediction of neo-epitope immunogenicity reveals TCR recognition determinants and provides insight into immunoediting,” Cell Rep. Med., vol. 2, no. 2, p. 100194, Feb. 2021.

[21] B. A. Albert et al., “Deep neural networks predict class I major histocompatibility complex epitope presentation and transfer learn neoepitope immunogenicity,” *Nat*. Mach. Intell., vol. 5, no. 8, pp. 861–872, Aug. 2023.

[22] G. Wang et al., “TLimmuno2: predicting MHC class II antigen immunogenicity through transfer learning,” Brief. Bioinform., vol. 24, no. 3, p. bbad116, May 2023.

[23] S. K. Dhanda et al., “Predicting HLA CD4 immunogenicity in human populations,” Front. Immunol., vol. 9, p. 1369, Jun. 2018.

[24] Z. Sethna et al., “RNA neoantigen vaccines prime long-lived CD8+ T cells in pancreatic cancer,” Nature, vol. 639, no. 8056, pp. 1042–1051, Mar. 2025.

[25] K.-H. Lee, T. J. Sears, M. Zanetti, and H. Carter, “NeoPrecis: enhancing immunotherapy response prediction through integration of qualified immunogenicity and clonality-aware neoantigen landscapes,” Nat. Commun., vol. 17, no. 1, p. 1966, Jan. 2026.

[26] N. Niknafs et al., “Persistent mutation burden drives sustained anti-tumor immune responses,” Nat. Med., vol. 29, no. 2, pp. 440–449, Feb. 2023.

[27] E. Hurtado, A. Bouchard-Côté, and A. Roth, “PhyClone: accurate Bayesian reconstruction of cancer phylogenies from bulk sequencing,” Bioinformatics, vol. 41, no. 7, p. btaf344, Jul. 2025.

[28] E. Lakatos et al., “Evolutionary dynamics of neoantigens in growing tumors,” Nat. Genet., vol. 52, no. 10, pp. 1057–1066, Oct. 2020.

[29] M. Perrinjaquet and C. Richard Schlegel, “Personalized neoantigen cancer vaccines: An analysis of the clinical and commercial potential of ongoing development programs,” Drug Discov. Today, vol. 28, no. 11, p. 103773, Nov. 2023.

[30] N. McGranahan et al., “Allele-Specific HLA Loss and Immune Escape in Lung Cancer Evolution,” Cell, vol. 171, no. 6, pp. 1259–1271.e11, Nov. 2017.

[31] L. Zapata et al., “Immune selection determines tumor antigenicity and influences response to checkpoint inhibitors,” Nat. Genet., vol. 55, no. 3, pp. 451–460, Mar. 2023.

[32] C. Puttick et al., “MHC Hammer reveals genetic and non-genetic HLA disruption in cancer evolution,” Nat. Genet., vol. 56, no. 10, pp. 2121–2131, Oct. 2024.

[33] A. Subramanian et al., “Personalized Circulating tumor DNA analysis for predicting outcomes and tracking response to radiotherapy and pembrolizumab in localized sarcomas: Analysis of the SU2C-SARC032 trial,” J. Clin. Oncol., no. JCO-26–00769, Sep. 2026, doi: 10.1200/jco-26-00769.

